# NAD^+^ Replenishment Reduces Cardiomyocyte Senescence and Improves Diastolic Function in the Aged Male Heart

**DOI:** 10.1101/2025.07.15.664959

**Authors:** Qingxia Huang, Yongjie Wang, Karina Farias, Xinliu Zeng, Qiuying Chen, Huiyong Cheng, Vijaya B. Baki, Steven S. Gross, Jacob Geri, Anthony A. Sauve, Yue Yang

**Author notes:** Deceased (Steven S. Gross, Anthony A. Sauve).

## Abstract

**Aims:** Cardiomyocyte senescence is a hallmark of cardiac aging and closely associated with diastolic dysfunction and heart failure in older adults. Strategies to effectively delay or reverse cardiomyocyte senescence in the aged heart remain limited. We investigated whether NAD⁺ replenishment regulates cardiomyocyte senescence and cardiac function in aging.

**Methods and Results:** Ventricular NAD⁺ metabolism, senescence markers, and cardiac function were assessed in young and aged mice of both sexes. Aged male mice exhibited reduced cardiac NAD⁺ levels due to impaired biosynthesis, limited mitochondrial mobilization, and increased consumption, leading to suppressed SIRT1 and SIRT6 activity and accumulation of senescent cardiomyocytes. These changes were accompanied by diastolic dysfunction, myocardial hypertrophy, and fibrosis. In cultured cardiomyocytes, NAD⁺ depletion induced cellular senescence, which was reversed by supplementation with the NAD⁺ precursors nicotinamide riboside (NR) and dihydronicotinamide riboside (NRH). In aged male mice, two months of NR or NRH treatment suppressed senescence markers, improved diastolic function, and reduced hypertrophy and fibrosis. Mechanistically, both NR and NRH activated SIRT1 and suppressed cell cycle arrest markers. NRH induced more robust SIRT1 activation, which deacetylated SIRT6, promoting its binding to histones and enhancing DNA damage repair. In contrast, aged female mice exhibited diastolic dysfunction without significant decline in cardiac NAD⁺ levels and did not respond to NR or NRH treatment.

**Conclusion:** NAD⁺ is a critical regulator of cardiomyocyte senescence in aging male hearts. NAD⁺ replenishment, particularly with NRH, reverses senescence-associated cardiac dysfunction through coordinated activation of SIRT1 and SIRT6. The sex-specific differences observed highlight distinct mechanisms underlying cardiac aging and emphasize the importance of considering biological sex in the development of NAD⁺- based therapeutic strategies.

**Translational Perspective:** Heart failure with preserved ejection fraction (HFpEF) disproportionately affects older adults and lacks effective therapies. This study establishes cardiomyocyte senescence as a therapeutically targetable mechanism in age-related diastolic dysfunction and positions NAD⁺ replenishment as a senescence-modifying intervention. By demonstrating that NAD⁺ precursors reverse senescence through coordinated SIRT1-SIRT6 activation, linking NAD⁺ metabolism to chromatin regulation and DNA damage repair, this work provides mechanistic insight into cardiac aging beyond metabolic effects alone. These findings support the clinical development of NAD⁺-based therapies for HFpEF and other age-associated cardiac conditions, with particular emphasis on patient stratification by sex and NAD⁺ status.

## 1. Introduction

Cardiomyocytes (CMs) make up the majority heart mass and are the primary cell type responsible for normal contractile function^1, 2^. With aging and prolonged stress, mitochondrial dysfunction, reactive oxygen species (ROS), and DNA damage response (DDR), these postmitotic and terminally differentiated cells can develop senescence-like phenotypes similar as those reported in proliferative cells^1, 3^. Senescent CMs exhibit canonical senescence markers, including elevated cell cycle inhibitors p16^INK4A^ and p21^WAF1/CIP1^, persistent DNA damage on telomeres, and increased senescence-associated beta-galactosidase (SA-β gal) activities from lysosome^4–6^. These CMs also secrete specific senescence-associated secretory phenotype (SASP) factors that can induce hypertrophy in adjacent CMs and fibrosis in cardiac fibroblasts, contributing to myocardial stiffness and both systolic and diastolic dysfunctions^4–7^. Increased senescent CMs were reported in aged mice^4, 8^, and in aged patients with heart failure with preserved ejection fraction (HFpEF)^4, 9^, the most common heart failure seen in older adults^3^. Targeted removal of senescent CMs, with the use of senolytic compounds and genetic clearance, have shown promises in improving cardiac function and reducing heart failure burden in mouse models^4, 10^. However, a mechanism-based therapy capable of both delaying onset and reversing established senescence in CMs is desirable for broader and safer application in the growing aging population.

NAD⁺ is one of the most abundant energy-sensing metabolites, functioning as a redox cofactor in hundreds of metabolic reactions and as a substrate for enzymes such as sirtuins (SIRTs), poly(ADP-ribose) polymerases (PARPs), and NAD⁺ hydrolases^11^. NAD^+^ depletion has been reported in myocardial tissues from both humans and mice with HFpEF and aging^12–14^. Low NAD^+^ levels can exacerbate mitochondrial dysfunction and cellular senescence, partially through reducing SIRT activities^15, 16^. SIRTs are a family of NAD^+^-dependent deacetylases, with nuclear SIRTs playing key roles in regulating inflammation, DNA repair, and senescence^17^. SIRT1 modulates transcription of cell cycle inhibitors via p53-p21 and Rb-p16^18^ pathways, and also controls SASPs release^17^. Another nuclear isoform, SIRT6, more directly regulates chromatin remodeling, telomere maintenance, DDR and genomic stability^19^. Knockout of SIRT6 induces CM senescence and premature death in mice^20^, suggesting its essential function in cardiac aging. Additionally, mitochondrial SIRT3 supports mitochondrial biogenesis and energy metabolism, both of which decline in aging and HFpEF^21^.

While NAD⁺ elevation can activate SIRTs and represents a potential anti-senescent strategy, the use of NAD⁺ precursors to directly target CM senescence has not been evaluated. Recent studies have shown that NAD⁺ supplementation improves HFpEF phenotypes in mice^14, 22–25^. For example, nicotinamide (NAM) improved diastolic function through SIRT1 activation in 24-month-old C57BL/6J mice^14^, while nicotinamide riboside (NR) enhanced mitochondrial function and reduced CM hypertrophy via SIRT1 and/or SIRT3 activation in models of pressure overloaded and diabetes-induced heart failure^23,24^. Although senescence was not directly assessed in these studies, the observed reduction in CM hypertrophy and fibrosis may reflect a decrease in senescent CMs and their SASP secretion^4^. Together, these findings support the mechanistic link between NAD⁺ levels, SIRT activation and CM senescence. However, they also highlight a critical knowledge gap in the direct evaluation of NAD⁺ metabolism in the progression and treatment of aging-related CM senescence, which limits the development of anti-senescence therapy for cardiac aging.

To address this gap, we investigated how age-related disruption of NAD⁺ balance in the heart is associated with cardiac senescence and SIRT activity in aged mice with cardiac dysfunction. We also compared the therapeutic effects of dihydronicotinamide riboside (NRH), a novel and highly potent NAD⁺ precursor discovered in our lab^26, 27^, to those of the widely used precursor NR. By examining their differential regulation of SIRT isoforms and target substrates, we aimed to understand how varying potentials of NAD⁺ elevation influence anti-senescence efficacy in aging hearts.

## 2. Material and Methods

### 2.1 Animals and Ethical Statements

C57BL/6JN mice aged 2-4 months (young) and 20-24 months (aged) of both sexes were obtained from the National Institute on Aging colony and housed in a specific pathogen- free facility under a 12-hour light/dark cycle at 23 ± 2°C with ad libitum access to standard chow and water. Body weight and food intake were monitored weekly. Mice received intraperitoneal (IP) injections of sterile PBS or NR or NRH (250 mg/kg in PBS) three times per week for 8 weeks. At termination of experiments, mice were anesthetized with ketamine (100 mg/kg) and xylazine (20 mg/kg, IP injection; Sigma-Aldrich, St. Louis, MO, USA). Adequate depth of anesthesia was confirmed by loss of pedal reflex before euthanasia via cervical dislocation under deep anesthesia. Hearts and lungs were collected and weighed, tibia length was measured, and remaining tissues were snap-frozen in liquid nitrogen and stored at −80°C.

All animal procedures were approved by the Institutional Animal Care and Use Committee of Weill Cornell Medicine (protocol #2015-0007) and conducted in accordance with the NIH Guide for the Care and Use of Laboratory Animals.

### 2.2 Statistical Analysis

All *in vitro* experiments were performed with at least three biological replicates and repeated in a minimum of three independent experiments with comparable results. Data are presented as mean ± standard deviation (SD). For comparisons involving more than two groups, one-way analysis of variance (ANOVA) was used, followed by Tukey’s or Dunnett’s post hoc test as appropriate. Comparisons between two groups were analyzed using unpaired two-tailed Student’s t-test. Post hoc testing was performed only when the ANOVA F-test reached statistical significance (P < 0.05) and assumptions of homogeneity of variance were met. Statistical analyses were performed using GraphPad Prism version 8.0 (GraphPad Software, Boston, MA, USA). A two-sided P-value < 0.05 was considered statistically significant. For metabolomic and proteomic pathway enrichment analyses, multiple-comparison correction was applied using the Benjamini-Hochberg false discovery rate (FDR) method to determine pathway-level significance.

### 2.3 Data Availability

The data supporting the findings of this study are available from the corresponding author upon reasonable request.

## 3. Results

### 3.1 NAD⁺ homeostasis is disrupted in aged male hearts

NAD⁺ levels were reduced 38% in ventricles of 24-month-old male mice compared to young controls (**Fig. 1A**). Untargeted metabolomics revealed nicotinate and nicotinamide metabolism among the most significantly altered pathways (**Fig. 1B, S. Table 1**). Related metabolites including NADH and NADP⁺ were similarly reduced, though NAD⁺/NADH ratios remained unchanged. Levels of NAD⁺ precursors (NAM, NMN, NaMN) decreased while the degradation product MeNAM increased, indicating reduced synthesis and increased degradation of NAD⁺ in aged hearts (**Fig. 1A**).

**Figure 1.**
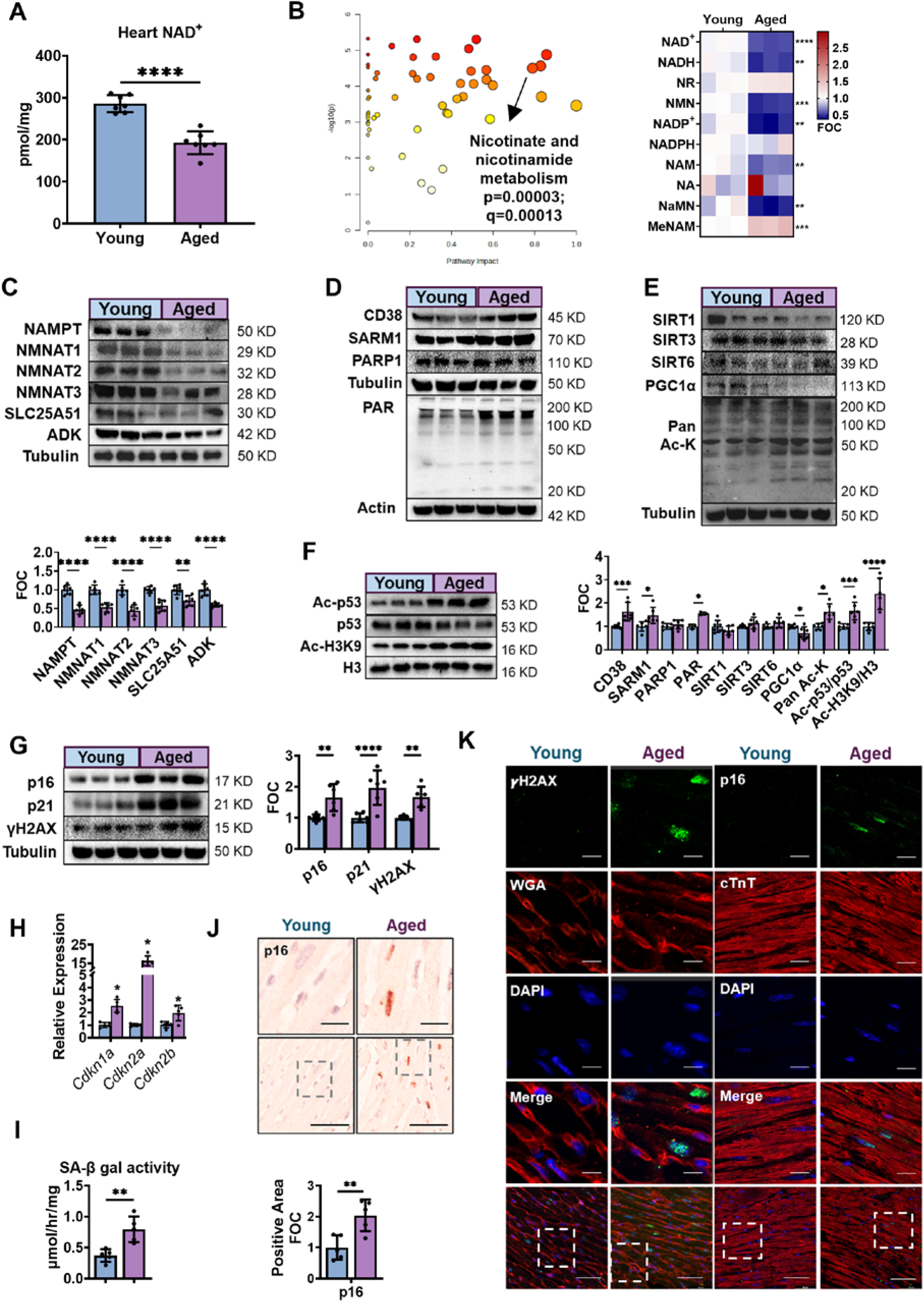
NAD^+^ imbalance accompanies increased cardiac senescence in aged hearts. **(A)** Ventricular NAD^+^ levels in young (2-4 month) and aged (24-month) male C57BL/6JN mice, N = 7 per group. **(B)** Untargeted metabolomic analysis revealed the top differentially impacted pathways between young and aged hearts (left). Heatmap on the right shows the relative abundance of metabolites as fold-over-control (FOC) in the Nicotinate and nicotinamide metabolism pathway, N = 3. **(C)** Representative western blot and densitometry quantification of NAD^+^ biosynthesis and mitochondrial storage enzymes in young and aged hearts, N = 6. **(D)** Representative expression of NAD^+^ consumers, along with the formation of poly-ADPR proteins (PAR) in the heart. **(E)** Expression of SIRT1, 3 and 6, PGC1α and pan acetyl-lysine (Ac-K) modification. **(F)** Immunoblots of SIRT1/6 targets Ac-p53 (K373/382) and Histone H3 (K9, H3K9), and the quantifications of **(D)**-**(F)**. N = 6 per group. **(G)** Expression of senescence markers in whole ventricle lysates, N = 6. **(H)** mRNA expression of cell cycle inhibitors between young and aged ventricles, N = 5. **(I)** Whole lysate SA-β gal activity measured by ONPG hydrolysis rate, N = 5. **(J)** Immunohistochemical DAB staining with Hematoxylin counterstaining and quantification of p16 in the ventricle sections, N = 5. Scale bars = 20 μm (upper) or 50 μm (lower). **(K)** Immunofluorescent staining of γH2AX/WGA/DAPI and p16/cTnT/DAPI to identify CM as the source of senescence upregulation. Scale bars = 20 μm (zoom in) or 50 μm. All data are shown in mean ± SD. Data were analyzed by unpaired t-test. *P < 0.05, **P < 0.01, ***P < 0.001 and **** P < 0.0001 in Aged vs. Young.

To identify the mechanisms underlying this NAD⁺ imbalance, we examined key biosynthetic and degradative enzymes. Biosynthetic enzymes NAMPT, NMNATs, and ADK^27^ were downregulated in aged ventricles (**Fig. 1C**). The mitochondrial NAD⁺ transporter SLC25A51 and NMNAT3 were also decreased, suggesting impaired mitochondrial NAD⁺ storage and mobilization^28, 29^. Conversely, NAD⁺ hydrolases CD38 and SARM1 were upregulated, consistent with previous studies identifying them as key contributors to age-related NAD⁺ decline^30–32^. PARP1 activity (assessed by PAR formation) was elevated despite unchanged protein levels (**Fig. 1D**). While SIRT1, SIRT3, and SIRT6 protein levels remained unchanged, aged hearts exhibited reduced deacetylase activity, evidenced by increased global lysine acetylation, reduced PGC-1α, and hyperacetylation of SIRT substrates p53-K382 and H3K9^33, 34^ (**Fig. 1E and F**). Together, NAD⁺ depletion results from reduced biosynthesis, impaired mitochondrial mobilization, and increased consumption by CD38, SARM1, and PARP1, consequently compromising SIRT activities

### 3.2 NAD⁺-deficient hearts exhibit elevated CM senescence

Since SIRT1 and SIRT6 play key roles in regulating senescence, we next examined senescence markers in aged hearts. Aged group exhibited elevated cell cycle arrest proteins p16 and p21, along with increased mRNA expression of *Cdkn1a*, *Cdkn2a*, and *Cdkn2b* (**Fig. 1G and H**; p16 antibody validation in **S. Fig. 1B**). The DNA damage marker γH2AX was also elevated, together with increased SA-β-gal activity (**Fig. 1G and I**), supporting the presence of a senescence phenotype.

To determine the cellular origin of these senescence signals, we performed immunohistochemical analysis. p16 immunostaining revealed signals confined to large, elongated nuclei characteristic of CM (**Fig. 1J**). Co-staining of γH2AX with WGA and p16 with cTnT confirmed that both markers localized to CM nuclei (**Fig. 1K**; quantification in **S. Fig. 1C**). Additionally, aged hearts showed increased expression of *Gdf15*, a CM-specific SASP factor implicated in promoting hypertrophy ^35^, as well as other SASP factors including *Tnfa*, *Il10*, and *Cxcl10* (**S. Fig. 1D**). These findings demonstrate that NAD⁺ deficiency is accompanied by marked cardiomyocyte senescence.

### 3.3 Aged mice exhibit diastolic dysfunction and mitochondrial impairment

We next examined how increased senescent CM burden impacts cardiac function. Aged male mice displayed common aging phenotypes including alopecia, whisker loss, and kyphosis. Echocardiography revealed slight but non-significant reductions in LVEF and FS, indicating preserved systolic function. In contrast, diastolic function was significantly impaired, as evidenced by elevated E/e′ ratios (**Fig. 2A; S. Table 2**). Aged mice also exhibited increased heart and lung weights, suggesting cardiac hypertrophy and pulmonary congestion, along with elevated serum pro-BNP levels (**Fig. 2B**). WGA staining confirmed CM hypertrophy, and Masson’s trichrome staining revealed increased interstitial and perivascular fibrosis (**Fig. 2C and D**). Collectively, these findings demonstrate that aged mice exhibit clinical features of aging-associated diastolic heart failure.

**Figure 2.**
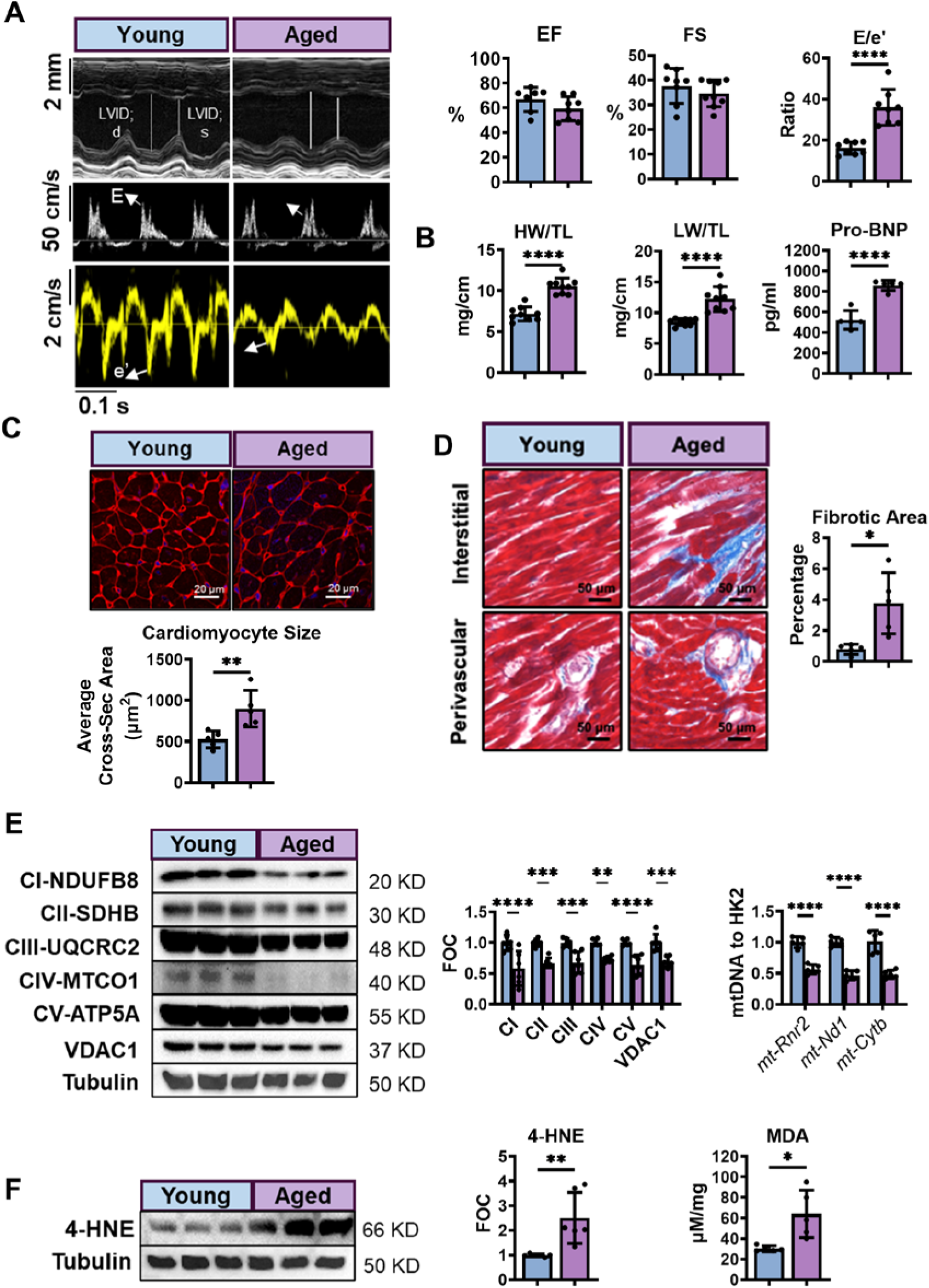
Mice with CM senescence exhibit aging-associated diastolic dysfunction with mitochondrial dysfunction. **(A)** Representative echocardiography-derived M-mode (top), pulsed-wave Doppler (middle) and tissue Doppler (bottom) tracings in young and aged male mice. Ejection fraction (EF) and fractional shortening (FS) percentages assess left ventricular systolic function, while the E/e’ ratios evaluate diastolic function. **(B)** The ratios of heart weight to tibia length (HW/TL), lung weight to tibia length (LW/TL) and the serum pro-BNP concentrations between young and aged mice. N = 7-8 for **(A)** & **(B)**. **(C)** WGA staining of the left ventricle slide (scale bars = 20 μm) and quantification of CM sizes. **(D)** Masson’s trichrome staining of left ventricular cross-sections (scale bars = 50 μm) and quantification of interstitial and perivascular fibrosis. **(E)** Mitochondrial quality examined by the protein expression and fold-over-control (FOC) of 5 OxPhos complexes and VDAC1, and the mitochondrial DNA (mtDNA) over nuclear DNA (HK2) ratios. **(F)** Presence of oxidative stress in heart lysate examined by immunoblot of lipid peroxidation product 4-HNE, and the quantification of the MDA activity assay between young and aged hearts. N = 5-6 per group. All data are shown in mean ± SD. *P < 0.05, **P < 0.01, ***P < 0.001 and **** P < 0.0001 in Aged vs. Young; data analyzed by unpaired t-test.

Mitochondrial dysfunction is a key characteristic of both diastolic heart failure and CM senescence^1, 36^. To assess mitochondrial integrity, we quantified OxPhos complex levels and found all five complexes were downregulated in aged hearts (**Fig. 2E**). VDAC1 was also reduced, along with decreased mtDNA copy number (**Fig. 2E**), indicating diminished mitochondrial quantity. Disruption of OxPhos can lead to ROS production and oxidative stress. Accordingly, lipid peroxidation markers 4-HNE and MDA were elevated in aged ventricles (**Fig. 2F**), indicating increased oxidative damage. Together, aged mice with increased senescent CM burden exhibit hallmark features of diastolic dysfunction accompanied by mitochondrial impairment and oxidative stress.

### 3.4 NAD⁺ availability determines cardiomyocyte senescence in vitro

To directly test whether NAD⁺ deficiency induces CM senescence, we treated AC16 cells, a human ventricular CM cell line, with the NAMPT inhibitor FK866 or the cardiotoxic agent DOX. FK866 reduced NAD⁺ levels to 50%, DOX to 30%, with combined treatment depleting to 20% (**Fig. 3A**). Inversely to NAD⁺ decline, SA-β-gal activity progressively increased (**Fig. 3B**), accompanied by elevated p16 protein levels (**Fig. 3C**), indicating a negative association between NAD⁺ availability and senescence induction. To confirm these effects in primary cells, we isolated CMs from young male mice and treated them with FK866. FK866 significantly reduced NAD⁺ levels (**Fig. 3D**) and increased expression of senescence markers p21 and γH2AX (**Fig. 3E**). Additionally, NAD⁺ depletion induced hyperacetylation of H3K9 (**Fig. 3E**), recapitulating the phenotypes observed in aged ventricular tissue and confirming that NAD⁺ deficiency alone can drive CM senescence.

**Figure 3.**
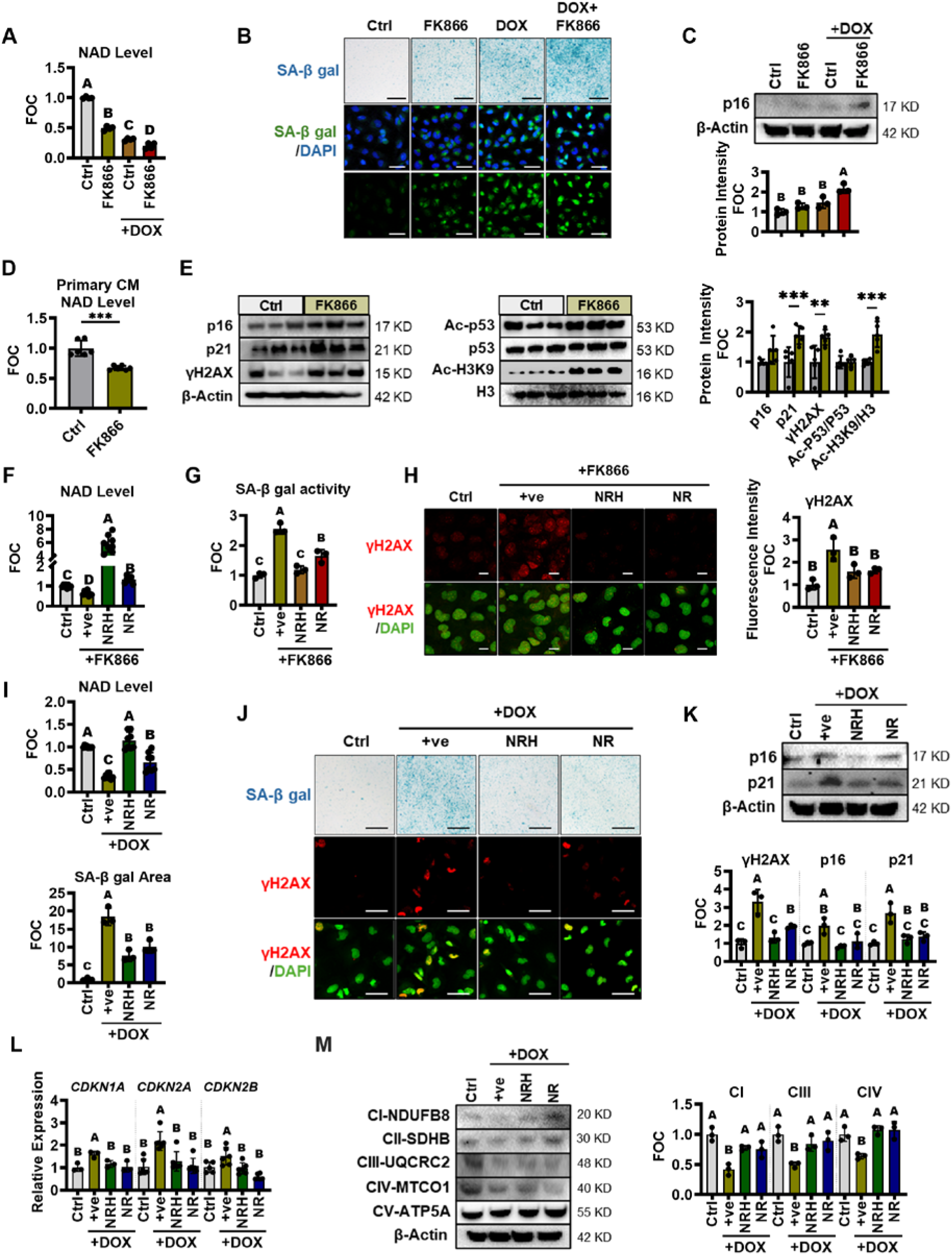
NAD^+^ availability drive *in vitro* CM senescence. **(A)** NAD^+^ levels in AC16 cells treated with 500 nM FK866 (NAMPT inhibitor) or/and 125 nM DOX for 48 hr expressed as fold-over-control (FOC). N = 4. **(B)** Representative image of SA-β gal staining in cells treated with FK866 or/and DOX, with blue color developed with X-Gal (scale bars = 200 μm) or fluorescent probe (scale bars = 50 μm). **(C)** The protein expression of p16 in FK866 or/and DOX treated cell lysates, N = 3. **(D)** NAD^+^ levels and **(E)** senescent markers, acetylated protein levels in primary CM isolated from young mice treated with 500 nM FK866 for 48 hr, N=5. **(F)** NAD^+^ levels in AC16 cells treated with 250 μM NRH or NR together with 500 nM FK866 for 48 hr. N = 8. **(G)** SA-β gal ONPG hydrolysis activities from cells treated with FK866 together with either NRH or NR for 48 hr, N = 3. **(H)** Representative IF staining of γH2AX with quantification. Scale bars = 50 μm, N = 3. **(I)** NAD^+^ levels (N=8) and quantified area of SA-β gal staining with X-Gal (N=3) in AC16 cells treated with 250 μM NRH or NR together with 125 nM DOX for 48 hr. **(J)** Representative images of SA-β gal staining shown in **(I)** (scale bars = 200 μm), and IF staining of γH2AX (scale bars = 50 μm). The immunoblots of p16 and p21, and the quantification of γH2AX, p16 and p21, N = 3. **(L)** The gene expression of cell cycle arrest markers. **(M)** Western blot and quantification of OxPhos Complexes. All data are shown in mean ± SD. Different letter (A, B or C) indicates statistically significant differences among groups (P < 0.05), as determined by ordinary one-way ANOVA followed by Tukey’s multiple comparisons test. Different letters denote groups that are significantly different from each other.

We next examined whether NAD⁺ restoration could reverse senescence. AC16 cells were treated with the NAD⁺ precursors NRH or NR (both at 250 μM) alongside FK866. While both precursors bypass NAMPT to enhance NAD⁺ synthesis, NRH increased NAD⁺ levels 5-fold compared to 1.5-fold by NR (**Fig. 3F**). Both attenuated FK866-induced SA-β-gal activity and γH2AX accumulation (**Fig. 3G and H**), indicating that NAD⁺ restoration rescues senescence induced by NAD⁺ deficiency.

Given that DOX induced pronounced NAD⁺ depletion and senescence, we tested how different doses of NRH and NR affect NAD⁺ restoration. Dose-response analysis showed NRH (100-250 μM) restored NAD⁺ more effectively than lower doses, while NR (500-1000 μM) plateaued at 250 μM without additional anti-senescent effects (**S. Fig. 2A and B**). At 250 μM, NRH restored NAD⁺ to near-control levels while NR only partially rescued DOX-induced depletion (**Fig. 3I**). Both significantly reduced SA-β-gal staining, γH2AX accumulation, and cell cycle arrest markers at protein and transcript levels (**Fig. 3J–M**). Unlike the senolytic agent ABT-263, which increased apoptosis, neither NAD⁺ precursor elevated apoptotic cell death (**S. Fig. 2C**), indicating senomorphic rather than senolytic effects. Moreover, both restored OxPhos complexes I, III, and IV that were downregulated by DOX (**Fig. 3N**). Together, these data demonstrate that CM NAD⁺ depletion drives senescence progression and that NAD⁺ replenishment effectively mitigates CM senescence.

### 3.5 NAD⁺ precursors utilize distinct metabolic pathways

The *in vitro* potency of NRH and NR as NAD⁺ boosters does not necessarily translate to the heart *in vivo*, where effectiveness depends on metabolic stability and delivery efficiency to CMs. To evaluate their *in vivo* potency, we measured ventricular NAD⁺ and related metabolites 30 minutes after acute IP injection. NRH induced a 2.3-fold increase in ventricular NAD⁺ levels versus 1.5-fold by NR (**S. Fig. 3A**). Metabolomic analysis revealed that NRH robustly increased nearly all NAD⁺-related metabolites, including 8-fold NRH itself, 5-fold NR, and 2.5-fold NMN (**S. Fig. 3B and C**). The elevation of NRH supports direct cardiac cell entry^37^, while increased NR suggests intracellular oxidation of NRH to NR. In contrast, NR treatment did not significantly increase NR or NMN levels but markedly elevated NaMN and NaAD, suggesting that nicotinic acid (NA), derived from NR via degradation to NAM followed by microbiome-mediated deamidation^38^, serves as the primary precursor for cardiac NAD⁺ synthesis. These data indicate that while both precursors elevate cardiac NAD⁺ levels *in vivo*, they are metabolized via distinct pathways.

### 3.6 NAD⁺ precursors improve diastolic function in aged male hearts

To determine whether long-term NAD⁺ precursor treatment reduces CM senescence and restores cardiac function, we randomized 22-24-month old male mice into treatment groups with matched baseline E/e′ ratios. Mice received 250 mg/kg NRH or NR by IP injection three times weekly for 8 weeks (**Fig. 4A**). Both treatments were well tolerated, with comparable body weight changes, blood glucose, and serum ALT levels (**S. Fig. 4A**). Since NRH lacks GRAS status, we further evaluated serum markers of liver and kidney function and observed no abnormalities (**S. Table 3**). Mortality rates were similar across control (3/10), NRH (1/10), and NR (2/8) groups (see **S. Methods**), indicating no adverse effects on survival.

**Figure 4.**
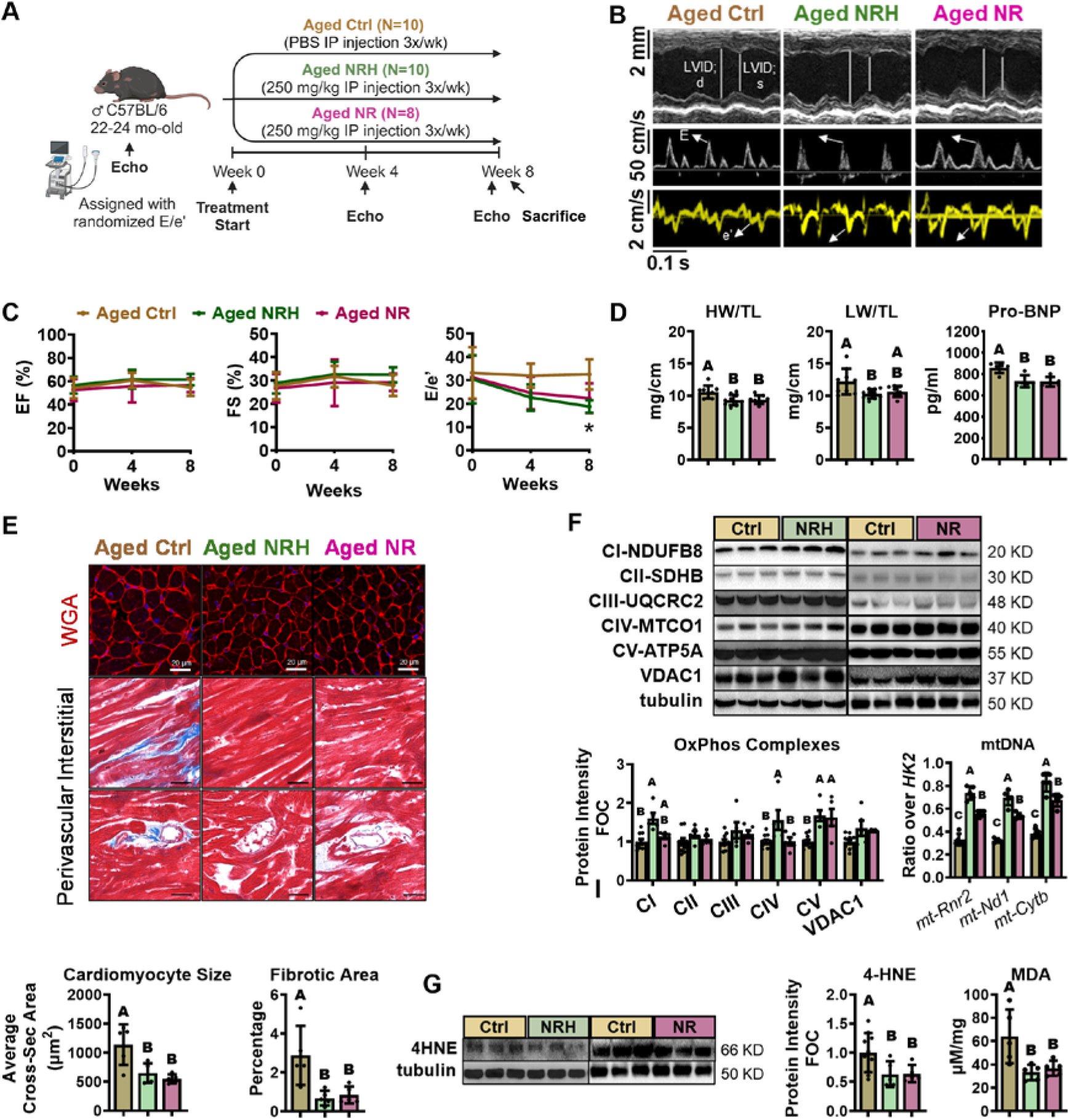
NAD^+^ precursor treatments lead to improved diastolic function and reduced cardiac senescence in aged male mice. **(A)** Aged male mice were randomly divided into 3 groups based on E/e’ ratio and subsequently administered 250 mg/kg NRH, NR, or PBS through IP injection 3 times a week for 8 weeks. **(B-C)** Representative echocardiography at week 8 and EF, FS and E/e’ ratios at week 0, 4 and 8, * indicate P < 0.05 between Ctrl and NRH group. **(D)** Heart weight and lung weight ratios, and serum pro-BNP levels by the end of 8 weeks. N = 6-10 for **(C)** & **(D)**. **(E)** Representative images and quantification of WGA-defined CM sizes (upper, scale bars = 20 μm) and Masson’s Trichrome stained cardiac fibrotic areas (lower, scale bars = 50 μm). **(F**) Representative levels of OxPhos Complexes quantified as fold-over-control (FOC) and mtDNA copy numbers in aged control and treated hearts. **(G)** Protein levels of 4-HNE and quantification of MDA activities in aged hearts. N = 5 for **(E) - (G)**. Data are shown in mean ± SD. Different letter (A, B or C) indicates statistically significant differences among groups (P < 0.05) and different letters denote groups that are significantly different from each other. Data were analyzed by ordinary one-way ANOVA followed by Dunnett’s multiple comparison test compared to Ctrl.

After 8 weeks, echocardiography revealed significantly reduced E/e′ ratios with NRH and a strong trend with NR (P=0.08), while LVEF and FS remained unchanged (**Fig. 4B and C; S. Table 4**), indicating improved diastolic function with preserved systolic function. Heart and lung weights decreased, serum proBNP levels lowered, and CM size and fibrotic area were reduced by both precursors (**Fig. 4D and E**), confirming attenuation of cardiac hypertrophy and fibrosis.

Mitochondrial function and antioxidant defense were also enhanced. NRH significantly increased OxPhos complexes I, IV, and V, whereas NR elevated only complex V (**Fig. 4F**). Consistent with this, NRH induced greater increases in mitochondrial genes *Rnr2*, *Nd1*, and *Cytb* compared to more modest elevations by NR (**Fig. 4F**). Both precursors significantly reduced oxidative stress markers 4-HNE and MDA (**Fig. 4G**), suggesting decreased ROS production and/or enhanced antioxidant capacity.

In contrast, aged (22-24-month) female mice showed mildly reduced cardiac NAD⁺ levels with greater inter-individual variability (**S. Fig. 4C**) and elevated E/e′ ratios (**S. Fig. 4D**) similar to males. However, neither NRH nor NR improved these parameters after 8 weeks (**S. Fig. 4E**), nor did they alter NAD⁺ metabolism enzymes (**S. Fig. 4F**), senescence markers (**S. Fig. 4G**), or OxPhos complexes, except for elevated 4-HNE levels (**S. Fig. 4H**). These findings suggest that cardiac dysfunction in aged female mice arises from distinct mechanisms and that NAD⁺ replenishment is most effective in hearts with pre-existing NAD⁺ deficiency and senescence.

### 3.7 Reduced CM senescence accompanies reprogrammed NAD⁺ metabolism

We next evaluated the senescent burden in aged hearts following NAD⁺ precursor treatment. Whole ventricular lysates showed reduced SA-β-gal activity with both NRH and NR (**Fig. 5A**). Consistently, protein and mRNA expressions of p16 and p21 were significantly downregulated by both precursors (**Fig. 5B and C**), indicating senescence-reducing effects. However, the DNA damage marker γH2AX and the CM-specific SASP factor *Gdf15* were significantly suppressed only by NRH (**Fig. 5B and C**), suggesting potentially greater impact on CM senescence. NRH also suppressed *Il10*, while both reduced *Gadd45a* and *Cxcl10* (**S. Fig. 5B**). Immunostaining confirmed that both precursors reduced p16 signal in CM nuclei, with NRH more substantially reducing γH2AX (**Fig. 5D**), consistent with whole ventricular protein expression.

**Figure 5.**
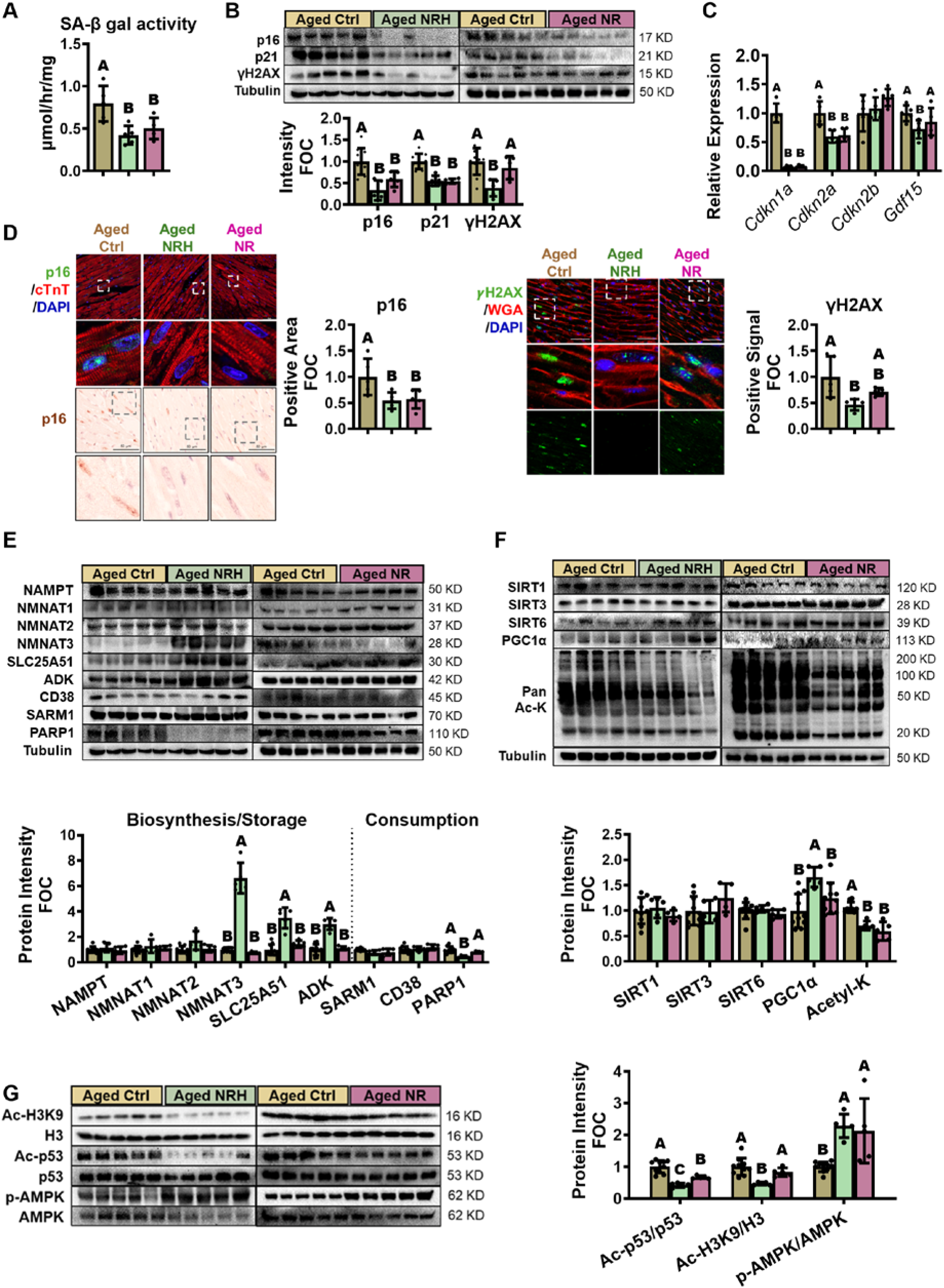
NAD^+^ precursor treatments suppress CM senescence and enhance SIRT activities in the aged male hearts. **(A)** Whole ventricular SA-β-gal activity determined by ONPG hydrolysis rate. **(B)** Representative western blots of senescent markers in ventricles and their quantification shown in fold-over-control (FOC). **(C)** mRNA expression of cell cycle inhibitors and CM specific SASP. **(D)** Representative immunofluorescent staining of p16/cTnT/DAPI and immunohistochemical staining of p16 with hematoxylin (left), and immunofluorescent staining of γH2AX/WGA/DAPI (right), along with their quantifications. Scale bars = 20 μm (zoom in) or 50 μm. **(E)** Western blots and quantification of NAD^+^ biosynthesis, storage and consumption-related enzymes. **(F)** SIRT-related proteins in the whole lysates. **(G)** Ventricular expressions of p53 and H3K9 acetylation, and AMPK phosphorylation, normalized to their total proteins. N = 5. All data are shown in mean ± SD. Different letter (A or B) indicates statistically significant differences among groups (P < 0.05), as determined by ordinary one-way ANOVA followed by Dunnett’s multiple comparison test compared to Ctrl (A) to (F) or Tukey’s multiple comparisons test (G). Different letters denote groups that are significantly different from each other.

Since NAD⁺ metabolism is significantly disrupted with aging, we examined whether precursor treatment influences not only acute NAD⁺ synthesis but also expression of key metabolic enzymes. While NR did not alter these components, NRH specifically upregulated the mitochondrial NAD⁺ transporter SLC25A51 and its partner NMNAT3 (**Fig. 5E**), as well as ADK. In terms of NAD⁺ consumption, NRH significantly reduced PARP1 protein levels (**Fig. 5E**), which may further decrease NAD⁺ depletion. These findings suggest long-term NRH supplementation enhances mitochondrial NAD⁺ storage and mobilization^29^, supporting sustained NAD⁺ replenishment to other subcellular compartments even after treatment cessation.

Given that the NAD⁺-senescence relationship is largely mediated by SIRTs, we evaluated key SIRT isoform expression and activity. While protein levels of SIRT1, SIRT3, and SIRT6 remained unchanged, global lysine acetylation was significantly reduced by both treatments and PGC-1α was upregulated by NRH (**Fig. 5F**), indicating enhanced SIRT activity. Both treatments led to p53 deacetylation, with NRH showing stronger effects (**Fig. 5G**). Since p53 acetylation enhances its stability and transcriptional activity, promoting expression of cell cycle arrest markers p16 and p21^39, 40^, the observed suppression of these markers supports SIRT1-mediated p53 deacetylation. In contrast, H3K9 acetylation, a primary SIRT6 target, decreased only with NRH. Since H3K9 acetylation is associated with chromatin relaxation and transcriptional activation^41^, its deacetylation by NRH likely contributes to further reduction of senescence-associated gene expression. Additionally, prior studies have suggested that NAD⁺ enhancement can paradoxically promote proinflammatory SASP secretion in senescent fibroblasts via AMPK inhibition^40^. Both precursors increased AMPK phosphorylation (**Fig. 5G**), indicating activation rather than inhibition in response to NAD⁺ elevation, potentially through the SIRT1-LKB1 pathway or altered AMP/ATP ratio^42^. Collectively, these findings suggest that while both NRH and NR activate SIRT1, only NRH also activates SIRT6, which may account for the greater anti-senescence efficacy observed with NRH in CMs.

### 3.8 NRH and NR induce distinct proteome deacetylation patterns

To identify specific SIRT-regulated targets underlying differential anti-senescence effects, we performed proteomic analysis on acetyl-lysine-enriched ventricular lysates. Principal component analysis revealed distinct profiles between aged controls and treated groups (**Fig. 6A**). NRH downregulated 167 proteins and NR downregulated 247 (fold change <0.75, P<0.05). Among 115 commonly suppressed proteins, cytoskeleton regulation was the most altered pathway, overlapping with focal adhesion, motor proteins, and muscle contraction (**Fig. 6B**), indicating a significant remodeling of CM structural components. From downregulated cytoskeleton proteins (**Fig. 6C**), we identified titin (TTN), a diastolic function regulator and SIRT1 deacetylation target^14^, and validated decreased TTN in acetyl-lysine immunoprecipitates (**Fig. 6D**). MYL4, implicated in hypertrophy with acetylation sites at K10 and K140^43^, also showed reduced acetylation (**Fig. 6D**), suggesting SIRT activation deacetylates filament proteins, contributing to reduced CM hypertrophy.

**Figure 6.**
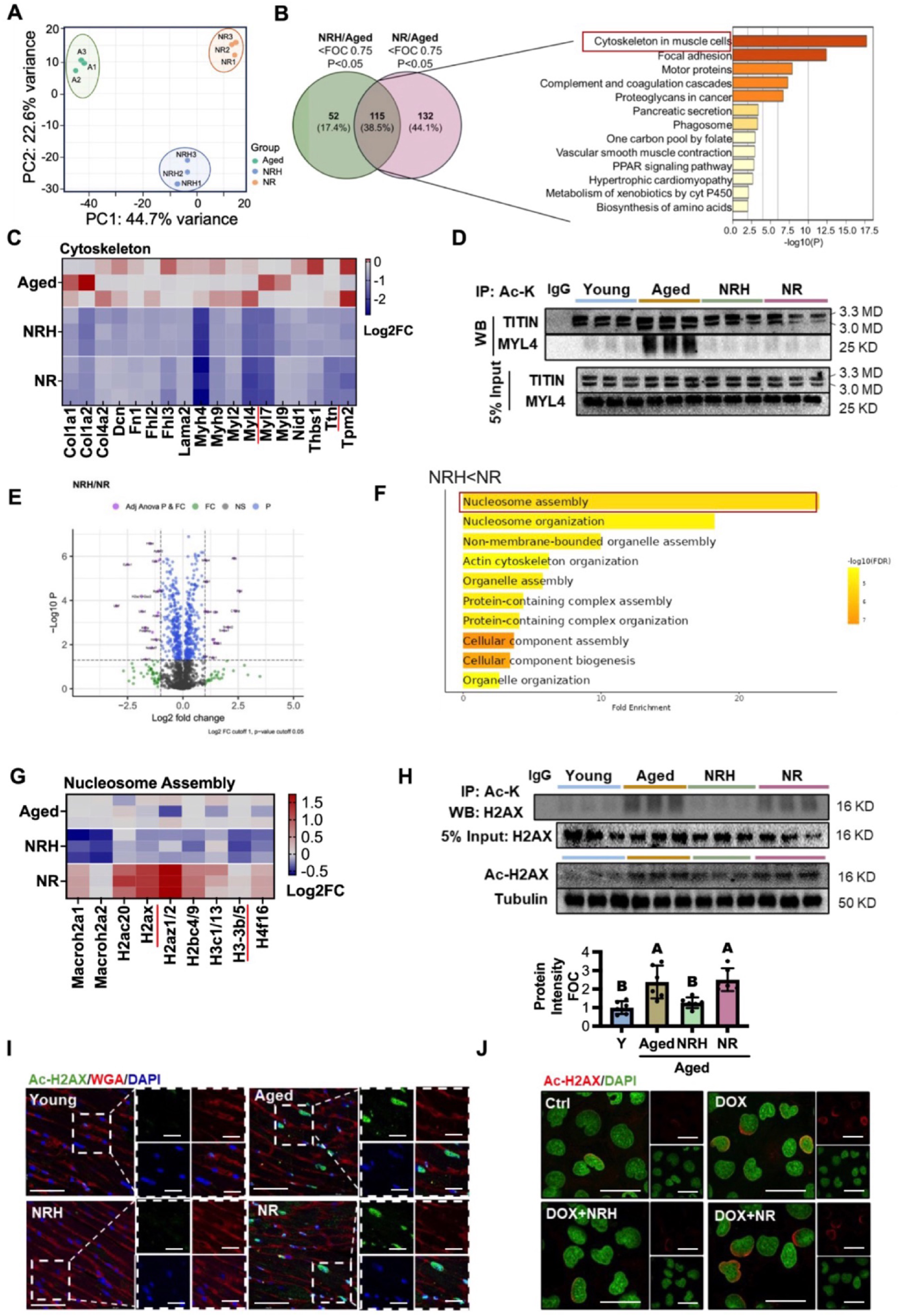
NAD^+^ precursors induce deacetylation in cardiac proteome. **(A)** Principal component analysis (PCA) of acetyl-lysine enriched proteome from Aged, NRH and NR treated hearts. **(B)** Venn diagram showing commonly downregulated proteins (FOC < 0.75, P < 0.05) in acetyl-lysine–enriched lysates from aged hearts treated with NRH or NR. KEGG pathways show the most significantly downregulated pathways by both NAD^+^ precursor treatments. **(C)** Heatmap showing the relative expression levels (Log2FC) of participants in the Cytoskeleton in muscle cells pathway. **(D)** Titin and MYL4 protein levels in acetyl-lysine immunoprecipitated lysates. **(E)** Volcano plot showing proteins differentially regulated by NRH and NR treatment-induced deacetylation on lysine. **(F)** The top 10 biological processes more significantly suppressed by NRH than NR (FOC < 0.75, P < 0.05). **(G)** Heatmap showing the expression levels of proteins participating in the Nucleosome assembly process. **(H)** H2AX expressions in acetyl-lysine immunoprecipitated lysates (upper) and immunoblot and quantification of H2AX acetylation at K-5 in whole heart lysates shown in fold-over-control (FOC) (lower). Different letter shows P < 0.05 between groups. N = 6. **(I)** IF images of Ac-H2AX in left ventricle, scale bar = 20 µm (zoom in) or 50 µm. **(J)** IF staining of Ac-H2AX in AC16 cells following treatment with NRH or NR, together with DOX for 48 hr, scale bar = 20 µm. All data are shown in mean ± SD, different letter shows P < 0.05 between groups. Data were analyzed with ordinary one-way ANOVA followed by Tukey’s multiple comparisons test. Different letter (A or B) indicates statistically significant differences among groups (P < 0.05) and different letters denote groups that are significantly different from each other.

Despite increased mitochondrial DNA and proteins, only 14 mitochondrial proteins were decreased by both treatments versus 57 were commonly increased, with TCA cycle as the top pathway (**S. Fig. 5A and B**). Upregulated proteins included TCA cycle, OxPhos, and fatty acid oxidation enzymes (**S. Fig. 5C**), reflecting increased abundance rather than decreased SIRT3 deacetylation. Supporting this, MSFragger analysis^44^ detected reduced acetylation of IDH2-K272 and ATP5PB-K225 despite unchanged total protein levels (**S. Fig. 5E**), indicating NAD⁺ replenishment activates SIRT3-mediated deacetylation, partially masked by increased mitochondrial protein abundance.

### 3.9 NRH specifically deacetylates histone proteins

Although NRH and NR shared many common targets, NRH uniquely deacetylated 53 proteins versus NR, with nucleosome assembly as the most differentially impacted process (**Fig. 6E and F**). Further analysis revealed nine histone proteins significantly deacetylated only by NRH (**Fig. 6G**), including H3.3, the most abundant H3 variant, supporting our observation that only NRH deacetylated H3K9. NRH-treated hearts showed reduced acetylation of H2A variants (H2AC20, H2AX, H2AZ) in immunoprecipitates, while NR-treated hearts showed slight increases (**Fig. 6G**), indicating major differences in histone regulation. H2AX contains acetylation sites at K5 and K37 associated with cell cycle arrest and DNA damage response^45^.

We validated that NRH but not NR reduced H2AX in acetyl-lysine immunoprecipitates (**Fig. 6H**). Similar patterns occurred for AcH2AX-K5 in whole lysates and ventricular immunofluorescence (**Fig. 6H and I**). In AC16 cells, DOX-increased H2AX acetylation was reversed by NRH but not NR (**Fig. 6J**). These findings demonstrate NRH shows superior efficacy in modulating histone acetylation, particularly H2AX and H3K9, which may regulate chromatin accessibility and senescence in CMs.

### 3.10 Differential SIRT6 activation mediates distinct anti-senescent effects

Given that histones were among the most differentially regulated proteins, we hypothesized differential SIRT1 or SIRT6 activation underlies the observed differences. Using siRNA in DOX-induced senescent AC16 cells (**S. Fig. 6A and B**), we found NRH’s SA-β-gal-reducing effect required both SIRT1 and SIRT6, while NR relied primarily on SIRT1 with minimal SIRT6 dependence (**Fig. 7A**). SIRT inhibitors yielded similar results (**S. Fig. 6C**). Supporting this, NRH reduced both p53 and H3K9 acetylation dependent on both SIRTs, whereas NR deacetylated only p53 (**Fig. 7B**), indicating both signal through SIRT1 but diverge in SIRT6-dependent H3K9 deacetylation.

**Figure 7.**
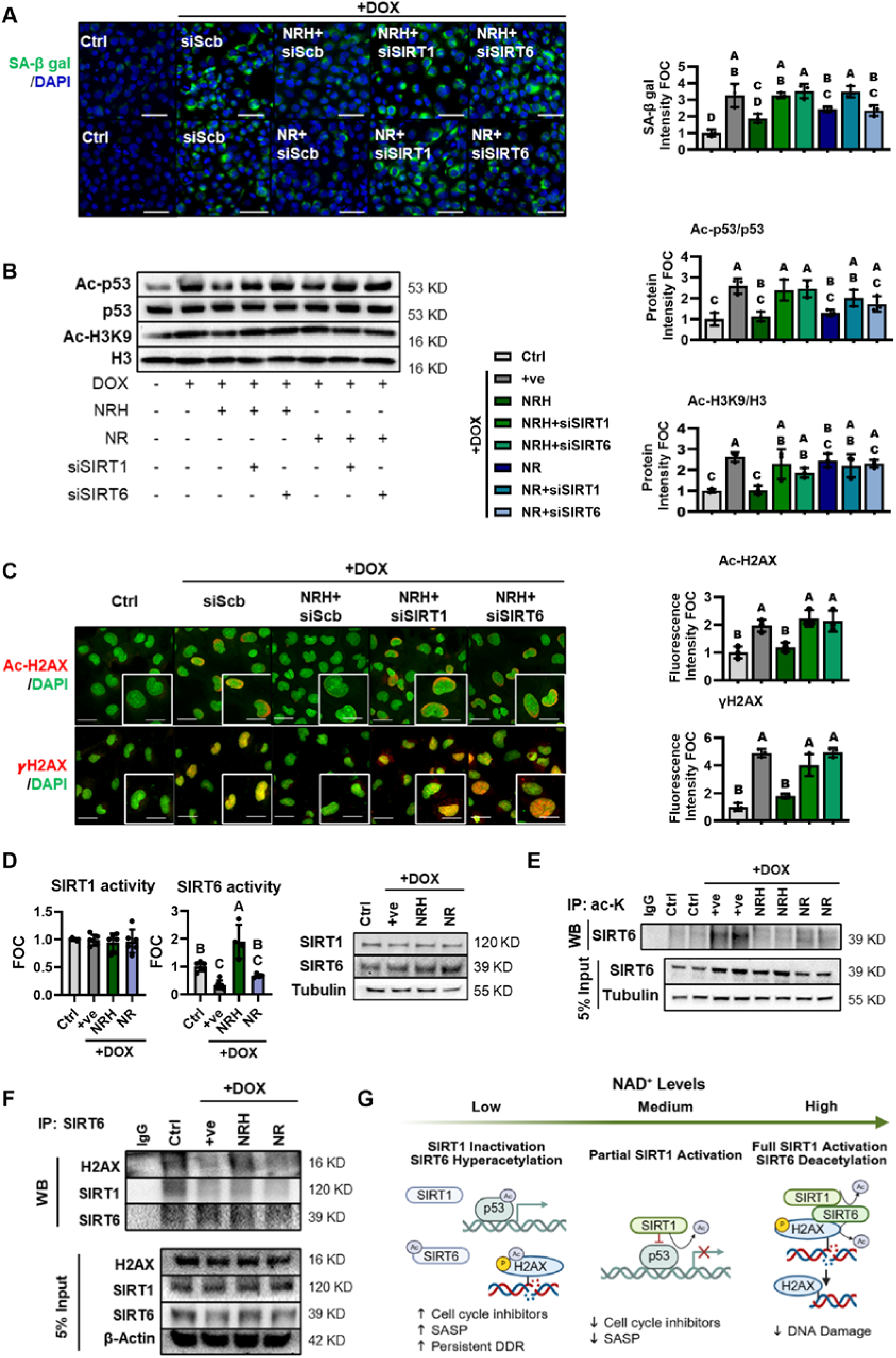
NRH specifically deacetylates H2AX through deacetylated SIRT6. **(A)** SA-β-gal staining with fluorescent probe in AC16 cells treated with DOX and NRH/NR, together with siRNA against SIRT1 or SIRT6 for 48 hr (scale bars = 50 μm), and their quantifications in fold-over-control (FOC). **(B)** Immunoblots of ac-p53/p53 and ac-H3K9/H3 (left), and their quantifications (right) in DOX-treated AC16 cells incubated with NAD^+^ precursors together with siRNA against SIRT1 or SIRT6. **(C)** IF staining and quantification of ac-H2AX and γH2AX expression in AC16 cells treated with DOX and NRH, with or without siRNA against SIRT1 or SIRT6. Scale bar = 20 µm or 10 µm (zoom in) **(D)** SIRT1 and SIRT6 activities measured in AC16 lysates treated with DOX ± 250 μM NRH or NR under saturated NAD⁺ conditions, along with corresponding SIRT1 and SIRT6 protein levels. **(E)** SIRT6 acetylation status examined in acetyl-lysine-immunoprecipitated lysates. **(F)** H2AX or SIRT1 expression in SIRT6-immunoprecipitated lysates in AC16 cells treated with DOX and NRH or NR. All data are shown in mean ± SD, N = 3. Different letter (A,B or C) indicates statistically significant differences among groups (P < 0.05), as analyzed by ordinary one-way ANOVA followed by Tukey’s multiple comparisons test. Different letters denote groups that are significantly different from each other. **(G)** Scheme of how NAD^+^ level changes in CMs alter SIRT1 and SIRT6 activities impacting on senescent pathways.

Focusing on H2AX, a prominently differential target, NRH suppressed H2AX acetylation (blocked by either SIRT knockdown), while NR had no effect (**Fig. 7C; S. Fig. 6D**). Since H2AX acetylation facilitates phosphorylation and DNA damage foci formation^45^, γH2AX levels correlated with acetylated H2AX, and NRH-mediated γH2AX reduction was blocked by either SIRT knockdown (**Fig. 7C**), suggesting H2AX deacetylation suppresses DNA damage and prolonged DNA damage response underlying senescence.

To determine whether differential NAD⁺ repletion by NRH and NR is the sole factor dictating SIRT1 and SIRT6 activation, we assessed SIRT enzymatic activities in AC16 lysates. Under saturated NAD⁺ concentrations with similar protein levels, SIRT1 activity was unchanged by either precursor (**Fig. 7D**) or by escalating doses of NRH (**S. Fig. 7A**), confirming the treatments did not alter SIRT1 expression or intrinsic activity but regulated SIRT1 activity through NAD⁺ concentration changes. In contrast, SIRT6 activity was reduced by DOX but restored only by NRH, even under saturated NAD⁺ (**Fig. 7D**), indicating NAD⁺-independent regulation. Lower NRH doses (50-100 μM) also failed to robustly activate SIRT6 (**S. Fig. 7B**) despite unchanged protein levels (**S. Fig. 7C**), suggesting SIRT6 activation requires both higher NAD⁺ repletion and functional regulation, potentially through post-translational modification.

Previous work showed SIRT1-mediated SIRT6 deacetylation promotes its recruitment to γH2AX for DNA repair^46^. Consistent with this mechanism, we found DOX increased SIRT6 acetylation, which was reduced by NRH but not NR (**Fig. 7E**). Co-immunoprecipitation revealed SIRT6 interactions with both H2AX and SIRT1 in controls were lost with DOX but restored by NRH, not NR (**Fig. 7F**), indicating SIRT6 deacetylation enhances its substrate affinity and enzymatic activity.

Collectively, we propose that distinct NAD⁺ precursors differentially activate SIRT pathways through graded NAD⁺ repletion. While moderate NAD⁺ increase by NR engages the SIRT1-p53 axis reducing cell cycle inhibitors and SASP factors, robust elevation by NRH promotes SIRT1-dependent SIRT6 deacetylation and activation, enhancing chromatin remodeling and DNA repair. This dual SIRT activation underlies NRH’s more pronounced anti-senescence effects in aged CMs (**Fig. 7G**).

## 4. Discussion

Aging is accompanied by accumulation of senescent CMs exhibiting cell cycle inhibitors, persistent DNA damage foci, and SASP, which are linked to myocardial dysfunction^4, 6^. Selective clearance of senescent CMs improves cardiac function, supporting the therapeutic relevance of targeting CM senescence^4, 6^. While upstream stressors such as telomeric DNA damage and mitochondrial dysfunction have been implicated^4, 47^, a clearly defined, targetable mechanism driving CM senescence has remained unclear. In this study, we identify NAD⁺ deficiency as a central, reversible driver of CM senescence. Aged hearts exhibited marked NAD⁺ depletion due to impaired biosynthesis, reduced mitochondrial mobilization, and accelerated degradation. *In vitro* NAD⁺ depletion was sufficient to induce senescence in AC16 cells and primary CMs. Mechanistically, reduced NAD⁺ suppressed SIRT1 and SIRT6 activity, causing hyperacetylation of transcription factors and histones, sustained DNA damage response signaling, and activation of senescence pathways. Restoration of NAD⁺ with NR and NRH improved diastolic dysfunction and reduced CM senescence, with NRH producing more robust effects. This work positions NAD⁺ restoration strategies as interventions that modify cellular senescence, with particular implications for aging-related diastolic dysfunction. Although prior studies have established beneficial effects of NAD⁺ supplementation in aging hearts^14, 22, 24^, our findings advance this field by revealing how NAD⁺ availability governs chromatin structure and DNA damage repair pathways, thereby deepening our mechanistic understanding of how metabolic and epigenetic networks converge to regulate cardiac aging.

A key novel finding is that the magnitude of NAD⁺ restoration determines selective activation of distinct SIRT pathways. The moderately potent precursor NR primarily activated SIRT1, reducing p53 activity and suppressing cell cycle arrest. In contrast, robust NAD⁺ elevation by NRH activated both SIRT1 and SIRT6, resulting in broader effects including reduced γH2AX accumulation and histone deacetylation. We identify H2AX as a functional SIRT6 substrate and demonstrate that SIRT6 activity is regulated by its acetylation status. Consistent with prior work showing SIRT1-mediated SIRT6 deacetylation promotes its recruitment to DNA damage sites^45,46^, our data support a SIRT1-SIRT6 axis only fully engaged when NAD⁺ levels are sufficiently restored. These findings highlight that Km-based predictions alone are insufficient to explain SIRT behavior in cells and emphasize the importance of acetylation status-mediated regulation (**Fig. 7G**).

From a therapeutic perspective, our data support NAD⁺ replenishment as a senomorphic strategy for aging-associated cardiac dysfunction. Unlike senolytic approaches that induce apoptosis and carry potential toxicity^7, 10^, NR and NRH suppressed CM senescence without increasing cell death. Moreover, despite reports of pro-inflammatory effects in other contexts^40,30^, we observed no inflammatory gene induction in the heart, suggesting favorable cardiac tolerance of NAD⁺ precursor treatment.

This study has several limitations that should be acknowledged. First, most of our cardiac analysis were not exclusively CM-specific, raising the possibility that signals from non-CM populations, such as fibroblasts, endothelial cells, or immune cells which can also undergo senescence^3^, may contribute to the observed effects. Although we confirmed the presence and modulation of senescence markers in CMs using immunofluorescence co-staining, more precise deconvolution approaches, such as CM-specific sorting or genetic labeling, will be required to definitively attribute ventricular-level changes to CMs.

Secondly, the aged female mice developed diastolic dysfunction without overt NAD⁺ deficiency and did not respond to NAD⁺ supplementation. While the underlying mechanisms remain unclear, this finding is consistent with prior reports demonstrating sex-specific differences in cardiac aging, with female C57BL/6 mice exhibiting relative resilience to adverse cardiac remodeling and dysfunction^48, 49^. In addition, late-life interventions have been shown to preferentially extend lifespan or confer functional benefits in male mice^50^. In our study, female hearts displayed greater heterogeneity in NAD⁺ levels, preserved expression of NAD⁺ biosynthetic enzymes, and reduced CM senescence burden, suggesting that sex-specific regulatory mechanisms may maintain NAD⁺ homeostasis and confer protection against CM senescence. This preserved NAD⁺ balance may also explain the limited responsiveness of female mice to precursor supplementation. Defining the molecular drivers of this sex-specific NAD⁺ regulation will be an important direction for future studies.

Finally, NAD⁺ precursors were administered via IP injection to achieve robust systemic exposure, which may limit direct translational relevance. Oral bioavailability of NAD⁺ precursors is known to be substantially lower^38^, yet our findings suggest that achieving sufficiently high tissue NAD⁺ levels is critical for therapeutic efficacy. These results highlight the potential value of developing more efficient, tissue-targeted NAD⁺ delivery strategies to maximize clinical benefit in cardiac aging.

In conclusion, this study identifies NAD⁺ homeostasis as a central regulator of CM senescence and diastolic dysfunction. We demonstrate that robust NAD⁺ replenishment, particularly with NRH, activates a cooperative SIRT1-SIRT6 program that promotes chromatin remodeling and DNA repair, resulting in a more pronounced anti-senescent effect. These findings provide mechanistic insight into how NAD⁺ metabolism governs cardiac aging and support targeting NAD⁺ pathways as a promising therapeutic strategy for age-associated cardiac dysfunction.

## Supporting information

Supplemental Methods, Tables, Figures and References

## Author Contribution

Q.H. collected the data, performed the analysis, wrote the paper; Y.W., K.F., X. Z., H.C. and V.B.B. collected the data; Q.C., S.S.G., J.G. provided consultant and resources for the analysis; A.A.S. obtained initial funding; Y.Y. conceived and designed the experiment, collected the data, wrote the paper and supervised the project. All authors discussed the results and contributed to the final manuscript.

