## Supplemental Methods, Tables, Figures and References for "NAD^+^ Replenishment Reduces Cardiomyocyte Senescence and Improves Diastolic Function in the Aged Male Heart"

#### Table of Supplemental Contents

### 1. Supplemental Methods

#### *1.1 Genotype verification, tissue processing, and animal adverse events*

C57BL/6JN genotypes were verified by PCR analysis of Nnt using genomic DNA according to [1], with amplification of both wild-type and mutant Nnt alleles. At the study endpoint, hearts and lungs were collected and weighed, and tibia length was recorded. Ventricular tissues were dissected, with samples from three mice per group used for protein extraction and immunoprecipitation, and samples from three to five mice per group fixed in 4% paraformaldehyde and embedded in paraffin for histological analyses. All remaining tissues were snap frozen and stored at −80 °C until analysis. During the study, 3/10, 1/10, and 2/8 mice in the control, NRH, and NR groups, respectively, were euthanized or died due to causes judged to be independent of treatment, including anesthesia-associated complications, rectal prolapse, or localized infection. Postmortem examination did not reveal any gross tumors in major organs.

#### *1.2 Chemicals*

NRH and NR used in this study were synthesized in-house, and their purity and integrity were determined by nuclear magnetic resonance (NMR) spectroscopy and high-performance liquid chromatography (HPLC) prior to use, as previously described{Zeng, 2025 #83}. All other chemicals were purchased from Sigma-Aldrich (St. Louis, MO, USA) unless otherwise specified.

#### *1.3 Echocardiographic Analysis*

Transthoracic echocardiography was performed using a Vevo 3100 imaging system (Fujifilm VisualSonics, Toronto, ON, Canada) equipped with a 30-MHz MS-400 transducer. Mice were lightly anesthetized with 1-2% isoflurane (Baxter Healthcare Corporation, Deerfield, IL, USA) delivered via nose cone and placed in the supine position on a temperature-controlled heating platform to maintain body temperature at 37°C. Transthoracic M-mode and B-mode views were recorded, and quantitative analysis was based on the average of three successive cardiac cycles under stable hemodynamic conditions. Left ventricular end-diastolic internal diameter (LVIDd) and left ventricular end-systolic internal diameter (LVIDs) were measured to calculate fractional shortening (FS) and ejection fraction (EF). The ratio between peak early-filling velocity of transmitral flow (E wave) and the corresponding mitral valve annulus velocity (e′ wave) was assessed using pulsed-wave Doppler and tissue Doppler imaging, respectively. All echocardiographic acquisitions and analyses were performed by the same experienced operator blinded to treatment groups and genotypes.

##### *1.4 Cell Culture*

Immortalized human ventricular myocytes (AC16 cells; ATCC, Manassas, VA, USA) were cultured in Dulbecco's Modified Eagle Medium/Nutrient Mixture F-12 (DMEM/F12; Thermo Fisher Scientific, Waltham, MA, USA) supplemented with 12.5% fetal bovine serum (FBS; Gibco, Thermo Fisher Scientific) and 100 U/mL penicillin-streptomycin (Gibco, Thermo Fisher Scientific) at 37°C in a humidified incubator under 5% $CO_2$. All senescence studies were performed using cells between passages 4 and 10.

##### *1.5 Isolation and culture of adult mouse cardiomyocytes*

Adult mouse cardiomyocytes were isolated using a Langendorff-free method as previously described[2]. Briefly, after anesthesia and thoracotomy, the heart was rapidly excised and immediately flushed through the right ventricle with EDTA-containing buffer. The ascending aorta was clamped, and enzymatic digestion was initiated by direct injection of collagenase solution into the left ventricle. The ventricular tissue was then minced and gently triturated to dissociate cardiomyocytes. The resulting cell suspension was filtered and subjected to four sequential rounds of gravity settling to enrich for viable cardiomyocytes. Rod-shaped, intact cardiomyocytes were counted and plated on laminin-coated culture dishes for subsequent experiments.

##### *1.6 Lentivirus infection*

Lentiviral particles carrying shRNA constructs targeting mouse p16 were generated by Genechem Co., Ltd. (Shanghai, China). Target sequence: GATGATGATGGGCAACGTTCA; control sequence: TTCTCCGAACGTGTCACGT. For knockdown experiments, cardiomyocytes at approximately 70% confluence were transduced with lentivirus at a multiplicity of infection (MOI) of 100 in the presence of HiTranG A transduction reagent. After 12 hours, the viral medium was replaced with fresh culture medium. Knockdown efficiency was subsequently validated by Western blot analysis.

##### *1.7 $NAD^+$ cycling assay*

Tissue and cellular $NAD^+$ concentrations were determined using our previously published cycling assay[3]. Briefly, $NAD^+$ was extracted with 7% perchloric acid and its concentration was determined in a cycling reaction with lactate, lactate dehydrogenase, diaphorase, and resazurin, enabling fluorescence quantification against an $NAD^+$ standard curve. Cellular concentrations were normalized to cell number, while tissue concentrations were normalized to tissue weight.

##### *1.8 Untargeted metabolomics*

For metabolite extraction, ~10 mg of snap frozen venticle tissues were pulverized in liquid nitrogen and extracted with 80% methanol. Remaining pellets were solubilized using 0.2 N NaOH for protein quantification and normalization. Untarget metabolomics was performed as described before[4]. Raw data files were processed using MassHunter Qualitative Analysis Software (B10.00; Agilent Technologies), with downstream comparative data analysis performed using MassHunter Profinder (B10.00) and MassProfiler Professional (Agilent, B15.1). Differentially impacted pathways were analyzed by MetaboAnalyst[5].

##### *1.9 Western blot*

To extract protein, cells or pulverized tissues were lysed with RIPA buffer containing protease inhibitor cocktail, phosphatase inhibitor, 5 μM Trichostatin A and 5 mM nicotinamide. Protein concentrations were determined by Bradford assay (Thermo Fisher Scientific). Equal amount of protein was separated by 10% SDS-PAGE gel and transferred to nitrocellulose or PVDF membranes, then incubated with primary antibodies (See Supplementary Table 5). Protein expression was visualized with HRP chemiluminescence (Thermo Fisher Scientific) and quantified using ChemiDoc™ imager (Bio-Rad, Hercules, CA, USA). All antibody information are listed in S. Table 5.

##### *1.10 Acetyl-lysine enriched proteomics*

For acetyl-lysine enriched proteomic analysis, ventricle protein lysates were immunoprecipitated with the Signal-Seeker™ Acetyl-Lysine Detection Kit, following the manufacturer's protocol (Cytoskeleton, Denver, CO). Proteomics profiling using label-free quantitation was performed by Orbitrap Fusion™ Lumos™ Tribrid™ Mass Spectrometer (Thermo Fisher). KEGG pathway analysis and Gene Ontology enrichment analysis were performed using DAVID[6] and ShinyGO[7]. Secondary analysis on post-transcriptional modified peptides were performed using MSFragger[8].

##### *1.11 Immunohistochemistry analysis*

Paraffin-embedded tissues were cut into 5 μm sections, then deparaffinized in xylene and rehydrated through a graded ethanol series. Antigen retrieval was performed by heating sections in citrate buffer (pH 6.0) at pressure cooker for 5 min. After blocking with 5% normal serum for 1 hr, sections were incubated with primary antibodies overnight at 4°C. Following treatment with HRP-conjugated secondary antibody for 1 hr at room temperature, DAB substrate was applied to detect HRP activity, and the nuclei were counterstained with hematoxylin. Images were taken using optical light microscope (Nikon). Masson's trichrome staining was performed to evaluate cardiac fibrosis according to the manufacturer's protocol (Diagnostic Biosystems). The percentage of the interstitial and peripheral fibrotic area in the left ventricle was calculated using the ImageJ software.

##### *1.12 Immunofluorescence staining*

AC16 cells were grown on glass coverslips and fixed in 4% paraformaldehyde for 10 min at room temperature and then permeabilized with 0.2% Triton X-100. The cell slides or heart sections after antigen retrieval were blocked with 5% BSA for 1 hr and then incubated with primary overnight at 4 °C. Then the samples were stained with Alexa Fluor 488-labeled secondary antibodies for 1 hr at room temperature in the dark. Nuclei were counterstained with DAPI, and slides were mounted with antifade medium. The images were taken using a Leica STELLARIS 5 confocal microscope system (Leica, Wetzlar, Germany).

To determine the cross-sectional area of cardiomyocytes, heart sections were stained with wheat germ agglutinin (WGA; AlexaFluorTM 594 conjugate; Invitrogen). Cardiomyocytes with a centrally located nucleus and a circularity index between 1.0 and 0.895 (radius ratio of 1:1 to 1:1.4) was measured. Three fields per sample were quantified for average cardiomyocyte size.

##### *1.13 Immunoprecipitation*

The immunoprecipitation assays were performed using protein A/G magnetic beads (L00277, GenScript). Briefly, AC16 cells were lysed in NP-40 lysis buffer supplemented with protease inhibitor cocktail. The cell lysates were then incubated overnight at 4°C with an anti-SIRT6 or anti-acetyl-lysine antibody (1:100), followed by a 1-hr incubation with magnetic beads at room temperature. After immunoprecipitation, the protein-beads complexes were washed, and the bound proteins were eluted by boiling in 30 μL of 1× loading buffer at 100°C for 5 min. The eluted samples were collected by magnetic separation, and the supernatant was stored for subsequent Western blot analysis.

##### *1.14 SA-β gal detection*

To colorimetric detection of senescent cells expressed a SA-β gal activity at pH 6.0, cells or tissues were fixed for immunostaining with X-gal following published protocol[9]. For fluorescent detection, fixed samples were incubated with CellEvent™ Senescence Green Probe (Invitrogen) and quantified using the Becton-Dickinson Celesta Analyzer

(BD Biosciences) or visualized under fluorescent microscope. Soluble SA-β gal activities in cell or tissue lysates were measured using 2-nitrophenyl-β-D-galactopyranoside (ONPG) in pH 6.0 cleavage solution[10]. Activity of SA-β gal was calulcated as nmoles ONPG hydrolyzed/hr/mg protein.

##### *1.15 Quantitative real-time PCR analysis*

Total RNA was extracted using Trizol (Thermo Fisher Scientific). Following quantification with a Nanodrop 2000c spectrophotometer (Thermo Fisher Scientific), cDNA was reverse-transcribed and synthesized with equal amounts of RNA using a commercial cDNA synthesis kit (ABclonal, MA, USA). Afterwards, quantitative PCR was performed on a QuantStudio 6 Flex Real-Time PCR system (Thermo Fisher Scientific) using QuantiTect SYBR Green PCR reagent (ABclonal) following manufacturer's instructions. The relative gene expression levels were calculated using the threshold cycle and the 2^(-ΔΔCT) method. β-Actin was set as a reference. All primer sequences used are shown in Supplementary Table 6 and Table 7.

##### *1.16 Mitochondrial DNA copy number determination*

DNA was isolated from tissues in lysis buffer with proteinase K at 55°C overnight following published protocol[11]. Mitochondrial DNA (mtDNA) copy number was quantified by qPCR using primers specific to the mitochondrial gene 16S, CytB and Nd1, and normalized to the nuclear gene HK2.

##### *1.17 N-Terminal Pro-B-Type Natriuretic Peptide (NT-proBNP) measurements*

NT-proBNP levels in serum were determined with a mouse ELISA kit (EK730273; AFG Bioscience) was used following the manufacturer's instruction.

##### *1.18 Thiobarbituric acid reactive substances (TBAR) assay*

To detect the lipid peroxidation product malondialdehyde (MDA), TBAR assay (Cayman, USA) was applied to equal amount of protein lysates following manufacturer's protocol. MDA concentrations were calculated using a standard curve and expressed as µM /mg protein.

##### *1.19 TITIN protein quantification*

To assess TITIN acetylation, protein lysates were subjected to immunoprecipitation with an anti-acetyl-lysine antibody, followed by separation on 4%-10% SDS-PAGE gels. Electrophoresis was performed at 50 V for 6-7 h, after which proteins were transferred to a nitrocellulose membrane in transfer buffer (10% methanol) at 30 mA overnight. The membranes were incubated with an anti-titin antibody (1:1000), and Tubulin was used as a loading control. Chemiluminescent signals were captured and quantified using a ChemiDoc™ imaging system (Bio-Rad, Hercules, CA, USA).

##### *1.20 siRNA transfection*

To knock down target genes, AC16 cells were transfected with Silencer™ Select Pre-Designed siRNA against human SIRT1 and SIRT6 (Invitrogen), or scramble siRNA, mixed with Lipofectamine 2000 for 48-72 hr.

##### *1.21 SIRT activity test*

SIRT1 and SIRT6 activities were measured with the FLUOR DE LYS® SIRT1 fluorometric drug discovery assay kit (Enzo Life Sciences, Inc.) and SIRT6 Fluorogenic Assay Kits (BPS Bioscience) respectively based on manufacturers' instructions. Briefly, equivalent amount of protein lysates (20 μg) from treated AC16 cells were incubated with fluorogenic, K382-acetylated p53 peptide (SIRT1) or K9-myristoylated Histone H3 peptide (SIRT6) lysine peptide substrates, saturated $NAD^+$ (5 mM) for 30 min at 37 °C, then incubated with SIRT developer containing 5 mM NAM for 15 min at room temperature. Activities were measured by fluorescence with excitation at 360 nm and emission at 460 nm, deducting negative control and expressed as percentage of control activity.

#### 2. Supplemental Tables

**Supplement Table 1. Differentially impacted metabolism pathways between young and aged mouse heart**

| KEGG Pathway | Raw P | FDR | Impact |
|---|---|---|---|
| Taurine and hypotaurine metabolism | 0.0000273 | 0.000133 | 0.829 |
| Nicotinate and nicotinamide metabolism | 0.0000315 | 0.000133 | 0.789 |
| Ascorbate and aldarate metabolism | 0.0000354 | 0.000139 | 0.568 |
| Purine metabolism | 0.0000381 | 0.000139 | 0.494 |
| Glycerolipid metabolism | 0.0000430 | 0.000144 | 0.044 |
| Pantothenate and CoA biosynthesis | 0.0000446 | 0.000144 | 0.214 |
| Amino sugar and nucleotide sugar metabolism | 0.0000560 | 0.000171 | 0.299 |
| Galactose metabolism | 0.0000610 | 0.000172 | 0.037 |
| Glyoxylate and dicarboxylate metabolism | 0.0000625 | 0.000172 | 0.235 |
| Pyrimidine metabolism | 0.0000658 | 0.000172 | 0.568 |
| Arginine and proline metabolism | 0.0000835 | 0.000205 | 0.444 |
| Citrate cycle (TCA cycle) | 0.0000859 | 0.000205 | 0.506 |
| Arginine biosynthesis | 0.0000940 | 0.000215 | 0.599 |
| Glycolysis / Gluconeogenesis | 0.0001939 | 0.000387 | 0.457 |

**Supplement Table 2. Echocardiographic parameters of young and aged mouse hearts**

| Parameter | Units | Young | Aged | P-Value |
|---|---|---|---|---|
| Group size | | 8 | 8 | |
| Heart Rate | BPM | 473.53±46.02 | 432.70±35.01 | 0.49 |
| GHDiameter;s | mm | 2.07±0.52 | 2.20±0.36 | 0.84 |
| Diameter;d | mm | 3.33±0.44 | 3.20±0.40 | 0.83 |
| Volume;s | uL | 15.38±10.55 | 16.86±6.31 | 0.90 |
| Volume;d | uL | 46.32±15.08 | 42.05±12.39 | 0.83 |
| Stroke Volume | uL | 30.93±6.56 | 25.19±6.49 | 0.55 |
| Ejection Fraction | % | 69.22±10.63 | 60.79±6.51 | 0.49 |
| Fractional Shortening | % | 38.44±7.82 | 31.68±4.64 | 0.47 |
| E | mm/s | 465.47±105.33 | 406.86±94.81 | 0.49 |
| e' | mm/s | 27.32±6.72 | 15.80±6.58 | 0.003 |
| E/e' | | 17.50±3.82 | 28.23±8.03 | 0.001 |
| Cardiac Output | mL/min | 14.50±2.66 | 10.88±2.81 | 0.37 |
| LVAW;s | mm | 1.96±0.58 | 1.58±0.30 | 0.57 |
| LVAW;d | mm | 1.47±0.50 | 1.24±0.13 | 0.66 |
| LVPW;s | mm | 1.72±0.43 | 1.39±0.33 | 0.55 |
| LVPW;d | mm | 1.38±0.55 | 1.12±0.37 | 0.70 |

**Supplement Table 3. Serum chemistry in Aged Ctrl and Aged NRH-treated mice**

| Parameter | Aged (N=5) | NRH (N=4) | P-Value |
|---|---|---|---|
| BUN (mg/dL) | 33.64±5.50 | 27.33±6.06 | 0.15 |
| CREA (mg/dL) | 0.13±0.03 | 0.29±0.38 | 0.38 |
| BUN/CREA Ratio | 260.42±34.84 | 208.42±122.94 | 0.39 |
| ALP (U/L) | 94.72±21.69 | 119.43±16.93 | 0.11 |
| ALT (U/L) | 21.10±7.37 | 24.93±4.90 | 0.40 |
| AST (U/L) | 89.84±63.46 | 78.00±43.47 | 0.76 |
| TBIL (mg/dL) | 0.14±0.09 | 0.15±0.06 | 0.93 |
| DBIL (mg/dL) | 0.01±0.004 | 0.02±0.005 | 0.34 |
| IBIL (mg/dL) | 0.13±0.09 | 0.13±0.05 | 0.97 |
| TP (g/dL)] | 4.87±0.19 | 4.64±0.35 | 0.23 |
| ALB (g/dL) | 2.99±0.07 | 2.82±0.19 | 0.09 |
| GLOB (g/dL) | 1.88±0.16 | 1.82±0.17 | 0.60 |
| A/G Ratio | 1.62±0.13 | 1.55±0.05 | 0.33 |
| P (mg/dL) | 6.72±1.16 | 6.52±0.36 | 0.76 |
| Ca (mg/dL) | 8.62±0.25 | 8.51±0.39 | 0.62 |
| GLU (mg/dL) | 133.50±16.89 | 162.18±32.33 | 0.13 |
| CHOL (mg/dL) | 86.22±12.24 | 98.15±4.44 | 0.11 |
| TRIG (mg/dL) | 52.68±3.76 | 47.08±12.18 | 0.36 |
| CK (U/L) | 370.00±776.72 | 141.08±170.76 | 0.59 |
| TCO2 (mEq/L) | 12.00±8.26 | 17.18±6.35 | 0.34 |
| Na (mEq/L) | 147.6±1.95 | 144.00±9.56 | 0.43 |
| K (mEq/L) | 5.16±1.36 | 5.33±1.35 | 0.86 |
| CL (mEq/L) | 109.80±1.79 | 107.25±7.04 | 0.45 |
| Na/K | 30.00±7.52 | 28.00±6.48 | 0.69 |
| Anion Gap | 31.20±6.72 | 25.00±9.06 | 0.28 |

**Supplement Table 4. Echocardiographic parameters at week 0 and 8**

| Parameter | Units | Ctrl Week 0 | Ctrl Week 8 | P-Value | NRH Week 0 | NRH Week 8 | P-Value | NR Week 0 | NR Week 8 | P-Value |
|---|---|---|---|---|---|---|---|---|---|---|
| Group Size | | 10 | 7 | | 10 | 9 | | 8 | 6 | |
| Heart Rate | BPM | 469.30 ±41.51 | 427.87 ±34.78 | 0.08 | 484.81 ±58.01 | 467.05 ±63.59 | 0.29 | 473.53 ±48.34 | 462.95 ±37.10 | 0.17 |
| Diameter;s | mm | 2.60 ±0.45 | 2.54 ±0.24 | 0.74 | 2.36 ±0.46 | 2.31 ±0.43 | 0.61 | 2.62 ±0.68 | 2.41 ±0.63 | 0.91 |
| Diameter;d | mm | 3.58 ±0.47 | 3.52 ±0.15 | 0.72 | 3.30 ±0.46 | 3.41 ±0.40 | 0.09 | 3.55 ±0.73 | 3.34 ±0.59 | 0.57 |
| Volume;s | uL | 25.79 ±10.08 | 23.49 ±5.63 | 0.60 | 20.57 ±9.40 | 19.43 ±8.88 | 0.69 | 27.67 ±18.40 | 22.4 ±10.89 | 0.92 |
| Volume;d | uL | 55.19 ±15.87 | 51.57 ±5.22 | 0.57 | 45.46 ±14.78 | 48.78 ±13.96 | 0.12 | 55.79 ±29.40 | 47.48 ±17.75 | 0.71 |
| Stroke Volume | uL | 29.40 ±8.70 | 28.08 ±3.12 | 0.71 | 24.89 ±5.47 | 29.34 ±5.97 | 0.01 | 28.12 ±11.93 | 25.08 ±8.68 | 0.30 |
| EF | % | 54.17 ±8.82 | 54.82 ±7.10 | 0.87 | 56.73 ±7.23 | 61.73 ±7.98 | 0.11 | 52.64 ±9.41 | 55.77 ±13.94 | 0.37 |
| FS | % | 27.63 ±5.83 | 27.92 ±4.55 | 0.93 | 28.98 ±4.60 | 32.59 ±5.49 | 0.08 | 26.65 ±6.11 | 29.06 ±9.96 | 0.38 |
| E | mm/s | 400.25 ±131.33 | 363.3 ±53.29 | 0.49 | 408.2 ±60.05 | 335.22 ±64.07 | 0.02 | 402.76 ±58.26 | 357.03 ±68.93 | 0.20 |
| e’ | mm/s | 12.27 ±2.25 | 11.32 ±1.21 | 0.33 | 15.09 ±7.15 | 16.35 ±4.68 | 0.66 | 13.83 ±4.02 | 16.59 ±4.09 | 0.23 |
| E/e’ | | 33.21 ±10.96 | 32.55 ±6.46 | 0.89 | 30.45 ±10.27 | 21.12 ±3.68 | 0.02 | 31.31 ±9.39 | 22.36 ±6.34 | 0.08 |
| | | 13.69 | 12.07 | 0.33 | 11.89 | 13.64 | 0.02 | 13.25 | 11.54 | 0.56 |

| | | | | | | | | | | |
|---|---|---|---|---|---|---|---|---|---|---|
| Cardiac Output | mL/min | ±3.86 | ±2.13 | | ±2.17 | ±3.05 | | ±5.54 | ±4.04 | |
| LVAW;s | mm | 1.83 ±0.41 | 1.71 ±0.30 | 0.49 | 1.72 ±0.31 | 1.66 ±0.34 | 0.32 | 1.72 ±0.25 | 1.8 ±0.48 | 0.92 |
| LVAW;d | mm | 1.51 ±0.44 | 1.37 ±0.32 | 0.47 | 1.38 ±0.33 | 1.31 ±0.37 | 0.60 | 1.42 ±0.30 | 1.48 ±0.50 | 0.37 |
| LVPW;s | mm | 1.59 ±0.45 | 0.98 ±0.14 | 0.003 | 1.85 ±0.52 | 1.5 ±0.43 | 0.12 | 1.52 ±0.53 | 1.37 ±0.42 | 0.08 |
| LVPW;d | mm | 1.37 ±0.45 | 0.73 ±0.07 | 0.002 | 1.64 ±0.54 | 1.25 ±0.46 | 0.05 | 1.32 ±0.48 | 1.19 ±0.33 | 0.04 |

**Supplement Table 5. Antibody information**

| Antibodies | Sourse | Identifier |
|---|---|---|
| NAMPT | IMGENE | IMG-6111A |
| NMNAT1 | RayBiotech | 144-61439 |
| NMNAT2 | NOVUS | 94693 |
| NMNAT3 | Thermo Fisher | PA5-113202 |
| SLC25A51 | Thermo Fisher | PA5-113407 |
| ADK | GeneTex | GTX101385 |
| CD38 | CST | 51000 |
| SARM1 | Thermo Fisher | PA5-20059 |
| PARP1 | CST | 9532 |
| SIRT1 | CST | 9475 |
| SIRT3 | CST | 2627 |
| SIRT6 | Abcam | ab191385 |
| PGC1α | CST | 2178 |
| Pan acetyl-lysine | CST | 9681 |
| Acetyl-Histone H3 (K9) | MilliporeSigma | ABE18 |
| Histone H3 | MilliporeSigma | H0164 |
| Acetyl-p53 (K382) | CST | 2525 |
| p53 | CST | 2524 |
| $p16^{INK4a}$ | Thermo Fisher | PA5-20379 |
| $p16^{INK4a}$ | CST | 80772 |
| $p21^{Cip1/Waf1}$ | CST | 37543 |
| γH2AX | CST | 9718 |
| Complex I-V | Thermo Fisher | 45-8099 |
| VDAC1 | Abcam | ab15895 |
| 4-HNE | Abcam | ab46545 |
| p-AMPK | CST | 2535 |
| AMPK | CST | 2532 |
| TITIN | Thermo Fisher | PA5-100211 |
| MYL4 | Thermo Fisher | PA5-103056 |
| H2AX | Thermo Fisher | PA1-41004 |
| Acetyl-H2AX (K5) | Thermo Fisher | BS-3781R |
| GAPDH | CST | 2118 |
| α/β-Tubulin | CST | 2148 |
| HRP-Rabbit IgG | CST | 7074 |
| HRP-Mouse IgG | CST | 7076 |

**Supplement Table 6. Mouse primer list**

| Name | Sequence |
| --- | --- |
| mt-Nd1-F | CTAGCAGAAACAAACCGGGC |
| mt-Nd1-R | CCGGCTGCGTATTCTACGTT |
| mt-Rnr2-F | CCGCAAGGGAAAGATGAAAGAC |
| mt-Rnr2-R | TCGTTTGGTTTCGGGGTTTC |
| HK2-F | GCCAGCCTCTCCTGATTTTAGTGT |
| HK2-R | GGGAACACAAAAGACCTCTTCTGG |
| mt-Cytb-F | CCCTAGCAATCGTTCACCTC |
| mt-Cytb-R | TGGGTCTCCTAGTATGTCTGG |
| Cdkn1a-F | CCGTGGACAGTGAGCAGTTGC |
| Cdkn1a-R | CCCTCCAGCGGCGTCTCC |
| Cdkn2a-F | TTCAGGTGATGATGATGGGCAACG |
| Cdkn2a-R | CGGGCGGGAGAAGGTAGTGG |
| Cdkn2b-F | TTGGGCGGCAGCAGTGAC |
| Cdkn2b-R | AGCGGTTCAGGGCGTTGG |
| Gadd45a-F | TGCTACTGGAGAACGACGC |
| Gadd45a-R | GGATCCTTCCATTGTGATGAA |
| Il10-F | GGACAACATACTGCTAACCGACTC |
| Il10-R | TGGATCATTTCCGATAAGGCTTGG |
| Cxcl10-F | TGCCTCATCCTGCTGGGTCTG |
| Cxcl10-R | CATTCTCACTGGCCCGTCATCG |
| Ccl5-F | GACACCACTCCCTGCTGCTTTG |
| Ccl5-R | CTCTGGGTTGGCACACACTTGG |
| Gdf15-F | TTGCTGCTGCTGCTGTCATGG |
| Gdf15-R | GCTCGTCGGCGTTGAGTTGG |
| TGFβ2-F | GAGCGGAGCGACGAGGAGTAC |
| TGFβ2-R | GAGCACCTGGGACTGTCTGGAG |
| TGFβ1-F | TGGCTCCTTCTCGGGCTCAC |
| TGFβ1-R | GGGACCCTGGGACACACTGG |
| Tnf-F | CCCTCACACTCAGATCATCTTCT |
| Tnf-R | GCTACGACGTGGGCTACAG |
| Il6-F | CTTCTTGGGACTGATGCTGGTGAC |
| Il6-R | AGGTCTGTTGGGAGTGGTATCCTC |
| Il1b-F | TCGCAGCAGCACATCAACAAG |
| Il1b-R | TCCACGGGAAAGACACAGGTAG |

**Supplement Table 7. Human primer list**

| Name | Sequence |
|---|---|
| CDKN1A-F | GTCACCGAGACACCACTGGAG |
| CDKN1A-R | CTGCCTCCTCCCAACTCATCC |
| CDKN2A-F | AGCAGCATGGAGCCTTCGG |
| CDKN2A-R | CGTAACTATTCGGTGCGTTGGG |
| CDKN2B-F | GGGAGGGCTTCCTGGACAC |
| CDKN2B-R | CGCTCCTCGGCCAAGTCC |

#### 3. Supplemental Figures

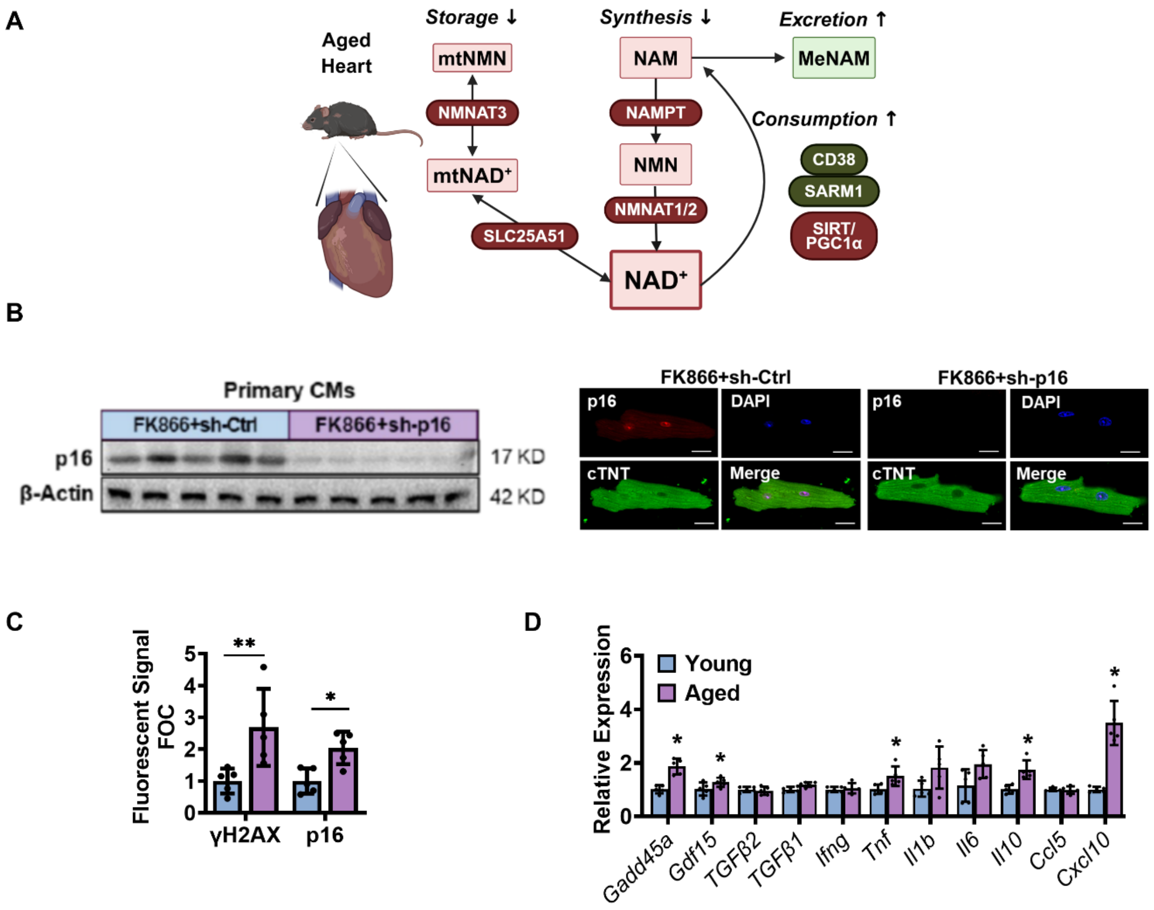

**Supplement Figure 1. NAD$^+$ metabolism and cardiac senescence in aged male mice. (A)** Illustration of mechanism leading to NAD$^+$ depletion in the aged heart. Downregulated metabolites and proteins are shown in red, and upregulated metabolites and proteins are shown in green. **(B)** Validation of p16 primary antibody in both immunoblotting and immunofluorescent imaging using primary CMs from young mice treated with FK866 and p16 shRNA. Scale bars = 20 μm. **(C)** Quantification of Fig.1K showing immunofluorescent staining of γH2AX or p16 positive nuclei in WGA or cTnT identified CMs. 5 fields were quantified and averaged for each animal, 5 animal slides were imaged. **(D)** mRNA expressions of SASP markers. N = 5. Data are shown in mean ± SD. * shows $P < 0.05$, **, $P < 0.01$. Unpaired t-test was used for statistical analysis.

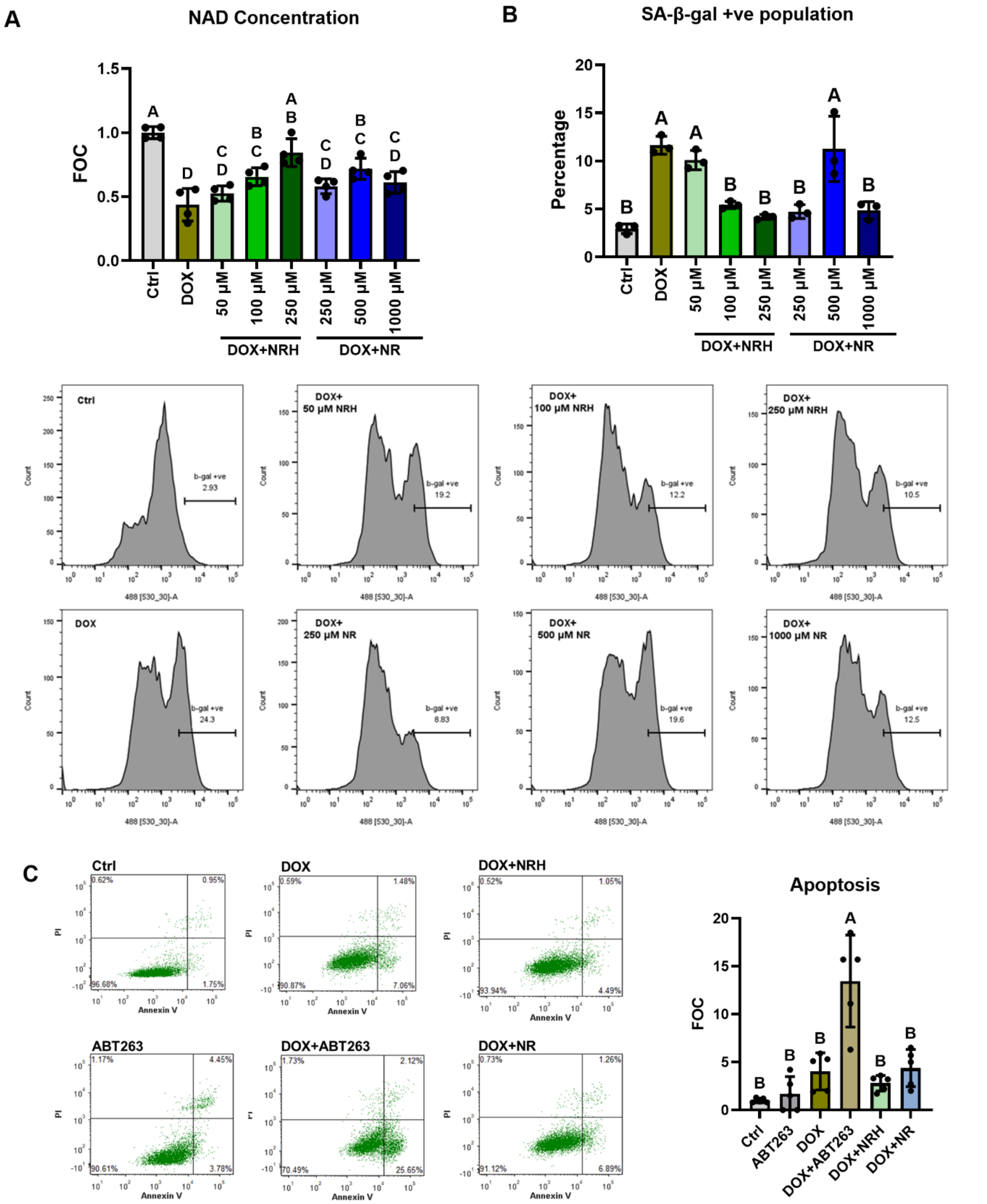

**Supplement Figure 2. Dosage response of NRH and NR in AC16 cells. (A)** Cellular NAD changes in DOX-treated AC16 cells co-incubated with different dosages of NRH or NR. N=4 per group. **(B)** Percentage of SA-β-gal positive cells in AC16 cells treated with DOX along with escalating dosages of NRH or NR for 48 hr determined with flow cytometry, and representative histograms. N=3 per group. **(C)** NRH or NR treatment at 250 μM did not induce apoptosis in DOX-treated AC16 cells compared to senolytic compound ABT263 at 2 μM, examined by Annexin V and PI staining, N=5 per group.

The bottom right quarter identified cells undergoing apoptosis, the bottom left were live cells, and the upper right were dead cells. Quantification of the bottom right quarter is shown on right. All data were analyzed by ordinary one-way ANOVA followed by Tukey's multiple comparisons test. Different letters identify significant difference ($P<0.05$) between groups, and same letters indicate no significant difference between groups.

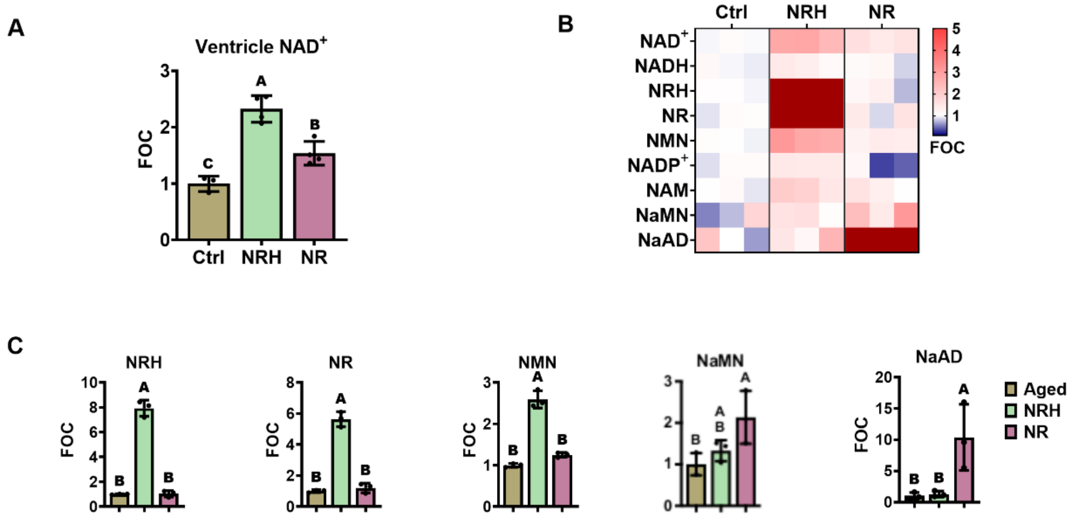

**Supplement Figure 3. NRH and NR enter cardiac cells via distinct metabolic pathways. (A)** Ventricle $NAD^+$ levels 30-min after IP injection of 250 mg/kg NRH or NR in young C57BL/6 mice. N = 4 per group. **(B-C)** Relative abundances of metabolites expressed as FOC in the Nicotinate and nicotinamide metabolism pathway extracted from the left ventricles 30-min post IP injection of NRH or NR. N = 3. Data shown as mean ± SD, different letters represent $P < 0.05$ when compared between groups. Data were analyzed by Tukey's multiple comparisons test following ordinary one-way ANOVA.

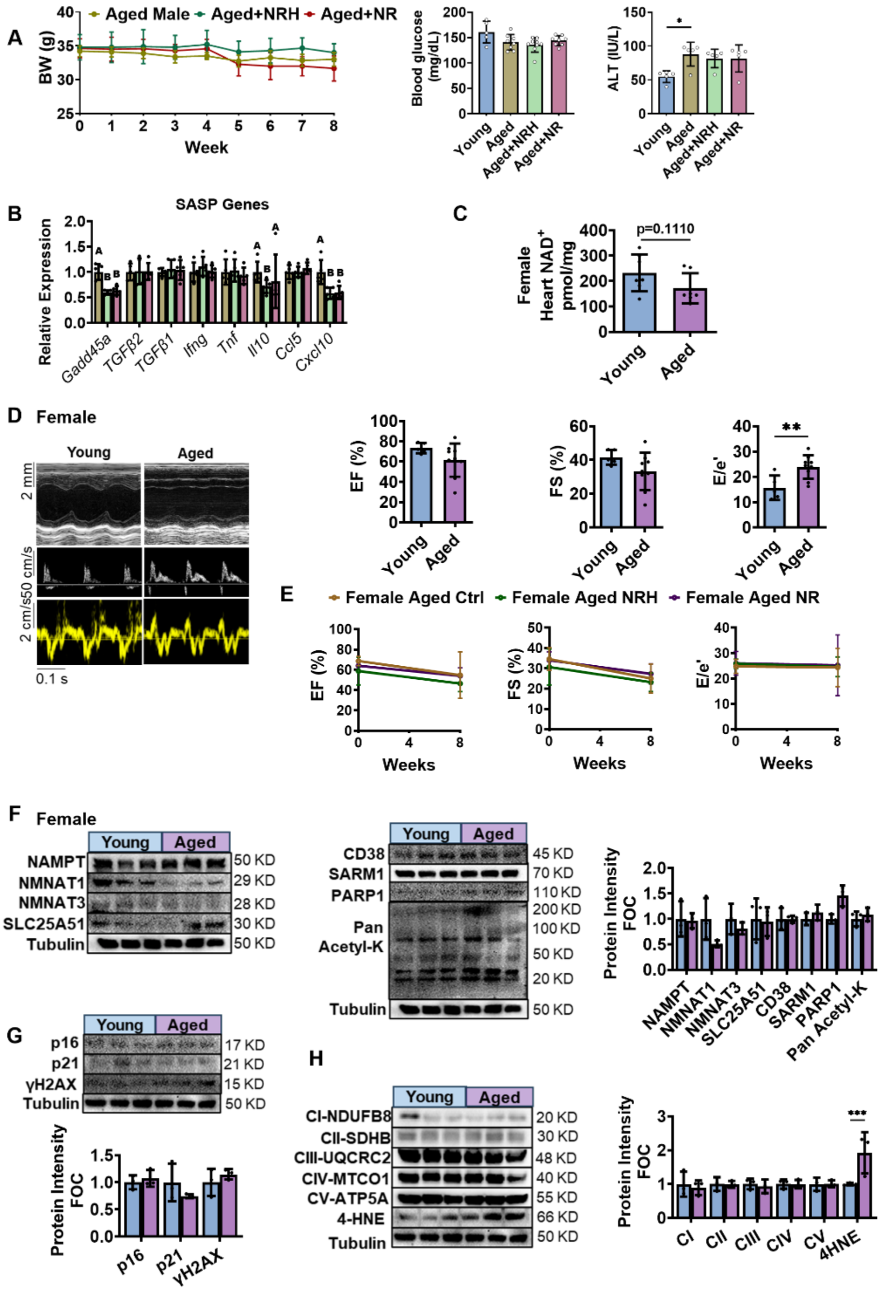

**Supplement Figure 4. Additional characterization of NAD+ precursor treatments in aged male and female mice. (A)** Body weight change throughout 8-week treatment and blood glucose and serum ALT levels at week 8 in aged male mice, N = 5-9 per group. **(B)** Additional SASP genes mRNA expression in aged male mice. N = 5 per group. For **(A)** and **(B)**, data were analyzed by ordinary one-way ANOVA followed by Tukey's multiple comparisons test. Different letters identify significant difference ($P<0.05$) between groups, and same letters indicate no significant difference between groups. **(C)** Cardiac NAD+ levels in young (3-4 month) and aged (22-24 month) female mice. **(D)** Representative echocardiography-derived M-mode (top), pulsed-wave Doppler (middle) and tissue Doppler (bottom) tracings in young and aged female C57BL/6 mice. EF and FS indicate left ventricular systolic function, and E/e′ indicate diastolic function. **(E)** Systolic and diastolic functions in aged female mice after 8 weeks of NRH or NR treatment. N = 7 per group for **(C-E)**. **(F)** Western blots and quantification of key $NAD^+$ biosynthesis/storage and consumption proteins in female hearts. **(G)** Expression and quantification of senescent markers in female hearts. **(H)** OxPhos complexes and 4-HNE expressions in young and aged female hearts. N = 3 for **(F-H)**. Data are shown in mean ± SD. * shows $P < 0.05$, **, $P < 0.01$, ***, $P < 0.001$ compared to young control. For **(C)** to **(H)**, data were analyzed by unpaired t-test.

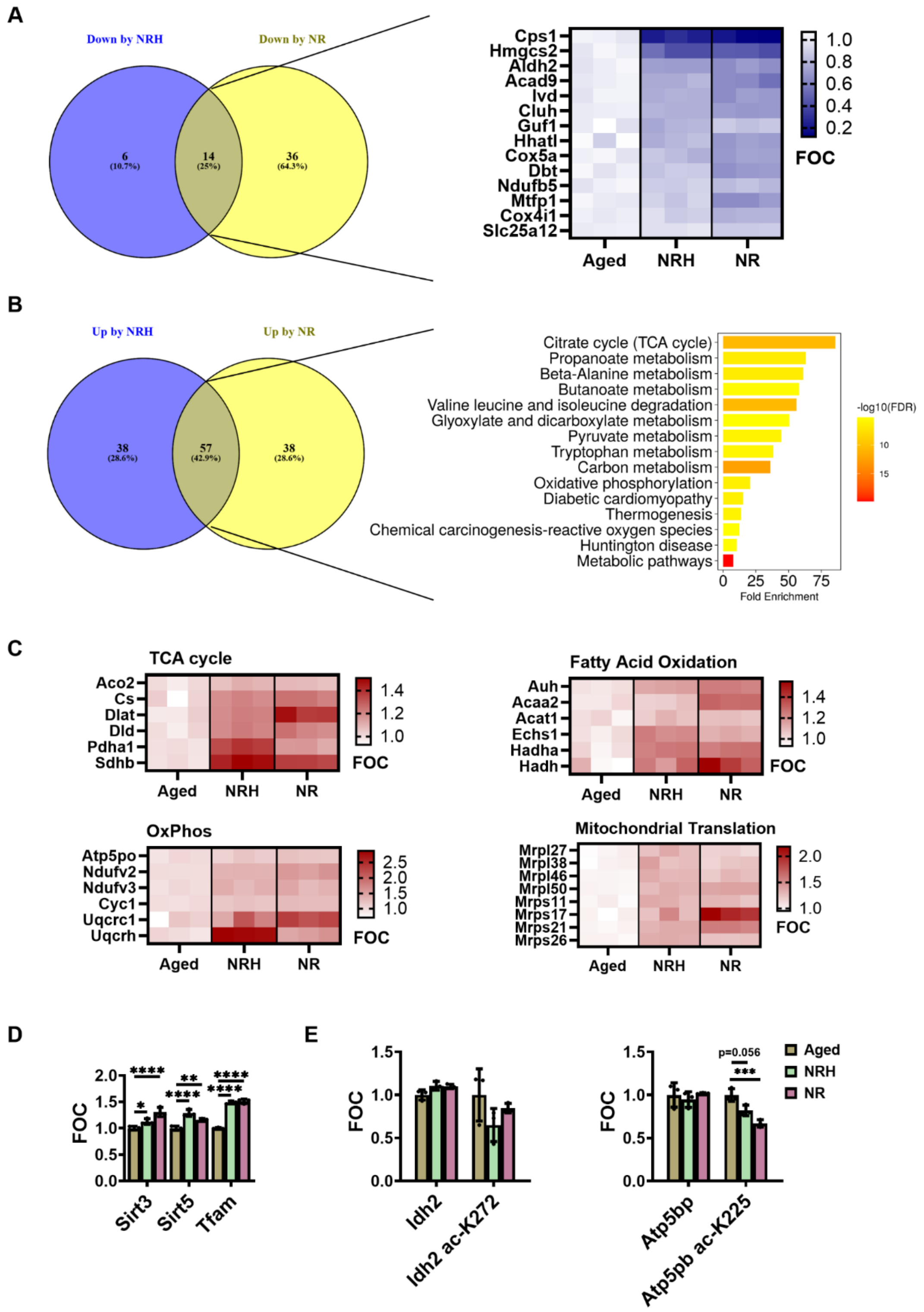

**Supplement Figure 5. Mitochondrial proteome differentially regulated by NAD$^+$ precursor treatments. (A)** Venn diagram and heatmap of commonly downregulated mitochondrial proteins by both NRH or NR treatment with $P < 0.05$ compared to Aged Ctrl. **(B)** Venn diagram and KEGG analysis showing the most enriched pathways in mitochondrial proteins commonly upregulated by both NRH and NR treatment compared to Aged Ctrl. **(C)** Heatmap showing components that are significantly upregulated with $P < 0.05$ by both NRH and NR in TCA cycle, OxPhos, Fatty acid oxidation and Mitochondrial translation pathways. **(D)** Relative levels of SIRT3, SIRT5 and TFAM in enriched proteome data. **(E)** MSFragger analysis identified acetyl-lysine modification on specific mitochondrial substrates. Data were analyzed by ordinary one-way ANOVA followed by Tukey's multiple comparisons test. Data were shown in mean ± SD, N = 3 per group. * shows $P < 0.05$, **, $P < 0.01$, ***, $P < 0.001$, ****, $P < 0.0001$.

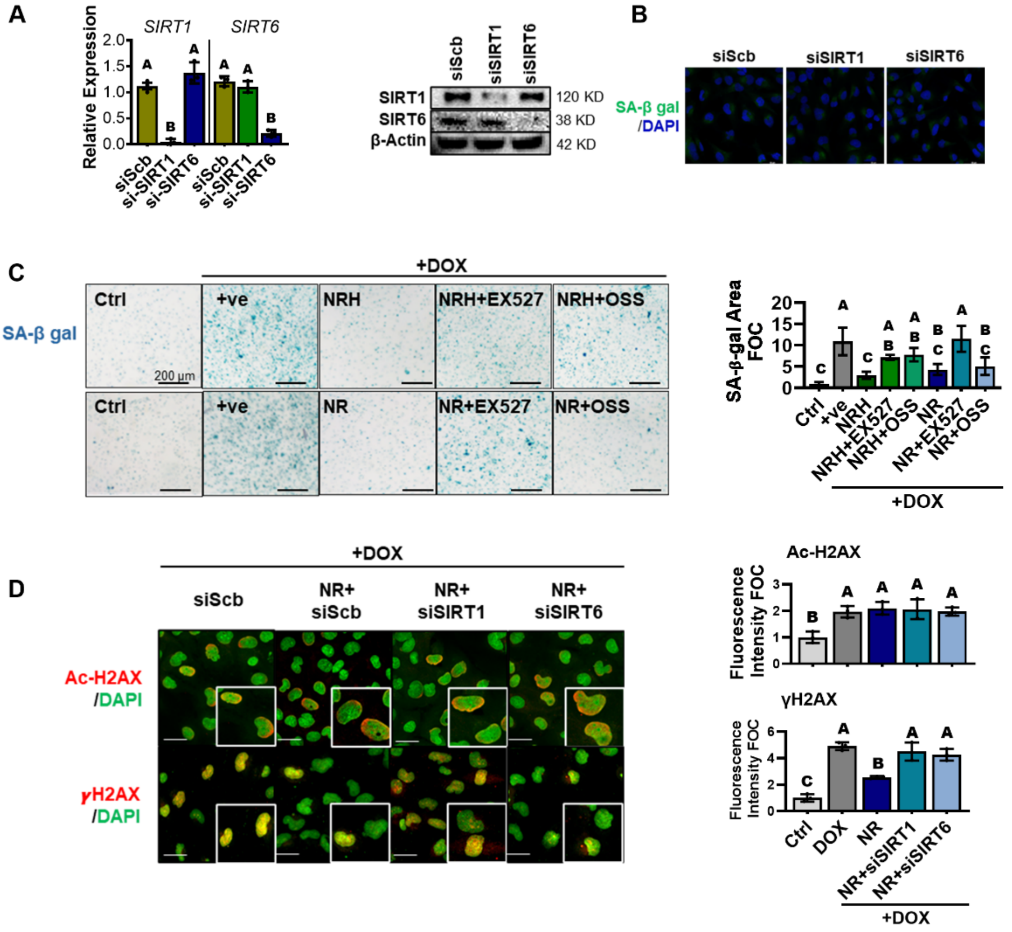

**Supplement Figure 6. Knockdown validation of si-SIRT1 or si-SIRT6 in AC16 cells. (A)** Left, mRNA expression of SIRT1 or SIRT6 in AC16 cells transfected with si-SIRT1, si-SIRT6 or siScramble (siScb) for 48 hr. Right, western blots of SIRT1 or SIRT6 in AC16 cells. **(B)** SA-β gal staining with fluorescent probe in AC16 cells treated with si-SIRT1 or si-SIRT6. **(C)** SA-β gal staining with X-gal in AC16 cells treated with DOX and NRH or NR, together with SIRT1 inhibitor (EX527) or SIRT6 inhibitor (OSS_128167), after 48 hr, and their quantification (right). **(D)** IF staining (scale bar = 20 μm) and quantification of ac-H2AX and γH2AX expression in AC16 cells treated with DOX and NR, with or without siRNA against SIRT1 or SIRT6. All data are shown in mean ± SD, N = 3. Data were analyzed by ordinary one-way ANOVA followed by Tukey's multiple comparisons test. Different letter shows $P < 0.05$ between groups.

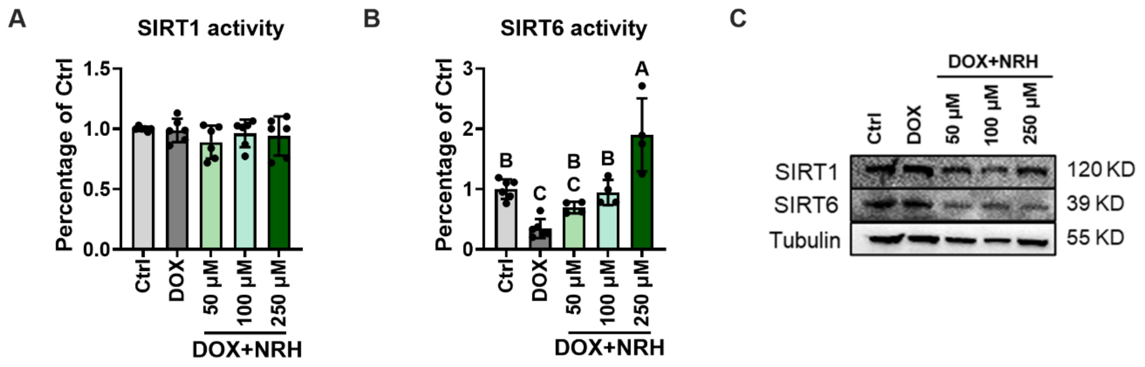

**Supplement Figure 7. SIRT1 and SIRT6 activity in different dosage of NRH. (A)** SIRT1 activity measured with equivalent amount of total cell lysate (20 μg) and saturated $NAD^+$ concentration (5 mM) in Ctrl, DOX treated cells co-incubated with escalating dosage of NRH for 48 hr. N=3 per group. **(B)** SIRT6 activity measured with equivalent amount of total cell lysate (20 μg) and saturated $NAD^+$ concentration (0.5 mM) in Ctrl, DOX treated cells co-incubated with escalating dosage of NRH for 48 hr. N=4-6 per group. **(C)** Immunoblotting of SIRT1 and SIRT6 protein levels in NRH treated cell lysates. All data are shown in mean ± SD. Data were analyzed by ordinary one-way ANOVA followed by Tukey's multiple comparisons test. Different letter shows $P < 0.05$ between groups.
